# Structure and dynamics of the ESX-5 type VII secretion system of *Mycobacterium tuberculosis*

**DOI:** 10.1101/2020.12.02.408906

**Authors:** Catalin M. Bunduc, Dirk Fahrenkamp, Jiri Wald, Roy Ummels, Wilbert Bitter, Edith N.G. Houben, Thomas C. Marlovits

## Abstract

*Mycobacterium tuberculosis* causes one of the most important infectious diseases in humans, leading to 1.5 million deaths every year. Specialized protein transport systems, called type VII secretion systems (T7SSs), are central for its virulence, but also crucial for nutrient and metabolite transport across the mycobacterial cell envelope. Here we present the first structure of an intact T7SS inner membrane complex of *M. tuberculosis*. We show how the 2.32 MDa, 165 transmembrane helices-containing ESX-5 assembly is restructured and stabilized as a trimer of dimers by the MycP_5_ protease. A trimer of MycP_5_ caps a central periplasmic dome-like chamber formed by three EccB_5_ dimers, with the proteolytic sites facing towards the cavity. This chamber suggests a central secretion and processing conduit. Complexes without MycP_5_ show disruption of the EccB_5_ periplasmic assembly and increased flexibility, highlighting the importance of this component for complex integrity. Beneath the EccB_5_-MycP_5_ chamber, dimers of the EccC_5_ ATPase assemble into three four-transmembrane helix bundles, which together seal the potential central secretion channel. Individual cytoplasmic EccC_5_ domains adopt two distinctive conformations, likely reflecting different secretion states. Our work suggests a novel mechanism of protein transport and provides a structural scaffold to aid drug development against the major human pathogen.

## Main

*Mycobacterium tuberculosis* encodes five homologous but functionally distinct type VII secretion systems (T7SSs), named ESX-1 to ESX-5. These systems translocate an arsenal of effector proteins across the unique and impermeable diderm cell envelope^1,2^. Because of their importance for mycobacterial physiology and virulence, T7SSs are considered promising targets for the development of novel tuberculosis drugs^3^. However, while T7SSs form hexameric complexes^4^, high-resolution structural information only exists for part of the T7SS, namely a dimeric ESX-3 subcomplex from the non-pathogenic strain *Mycobacterium smegmatis*^5,6^. In this study, we reconstituted the ESX-5 T7SS of *M. tuberculosis* H37Rv (ESX-5_mtb_) in *M. smegmatis* to obtain a structural view of the entire, hexameric T7SS membrane complex from this important human pathogen.

### Architecture and stoichiometry of an intact T7SS

The ESX-5_mtb_ system showed robust expression in *M. smegmatis* and proper assembly of the membrane complex, when produced from a low copy number plasmid and under control of native promoters (Extended Data Fig.1a, b). Excitingly, purification of the ESX-5_mtb_ membrane complex using a C-terminal StrepTag on EccC_5_ and mild solubilization conditions resulted in co-purifiction of the conserved MycP_5_ protease (Extended Data Fig. 1c, d, e, 3). This component was absent in all previously reported T7SS structures^4–6^, although MycP is known to be essential for T7SS functioning and complex stability^7^.

Cryo-EM analysis of the ESX-5_mtb_ complex, showed clear hexameric particles (Extended Data Fig. 4, 5). *Ab initio* reconstruction of the full complex without symmetry enforcement yielded an average resolution of ~4 Å (Extended Data Fig. 5, 6), which improved to an overall ~3.5 Å after recentering, focused classifications and signal subtractions, allowing to build ~78% of the stable complex *de novo* (Extended Data Table 2). Towards the periphery of the complex the resolution gradually decreases (Extended Data Fig. 6). The ESX-5_mtb_ complex is composed of six copies of EccB_5_, EccC_5_ and EccE_5_, twelve copies of EccD_5_ and three copies of MycP_5_ (Fig. 1b, c, d), yielding a 2.32 MDa complex that is anchored in the inner membrane (IM) via an astonishing number of 165 transmembrane helices (TMHs) (Fig. 1e). The membrane assembly can be best described as a trimer of dimers where each dimer is composed of two protomers of EccB_5_:EccC_5_:EccD_5_:EccE_5_ in a ratio of 1:1:2:1, and a single MycP_5_ copy (Fig. 1f). In the intact complex, the angle between protomers at the membrane level differs by ~0.5 degrees, with an angle of 59.7 degrees between protomers from one dimer to 60.2 degrees between protomers of adjacent dimers. However, the angles between protomers at cytosolic level differs by >10 degrees, as protomers from a dimer display an angle of 65.1 degrees, whereas two protomers from adjacent dimers form an angle of 54.8 degrees (Extended Data Fig. 8).

**Fig. 1.**
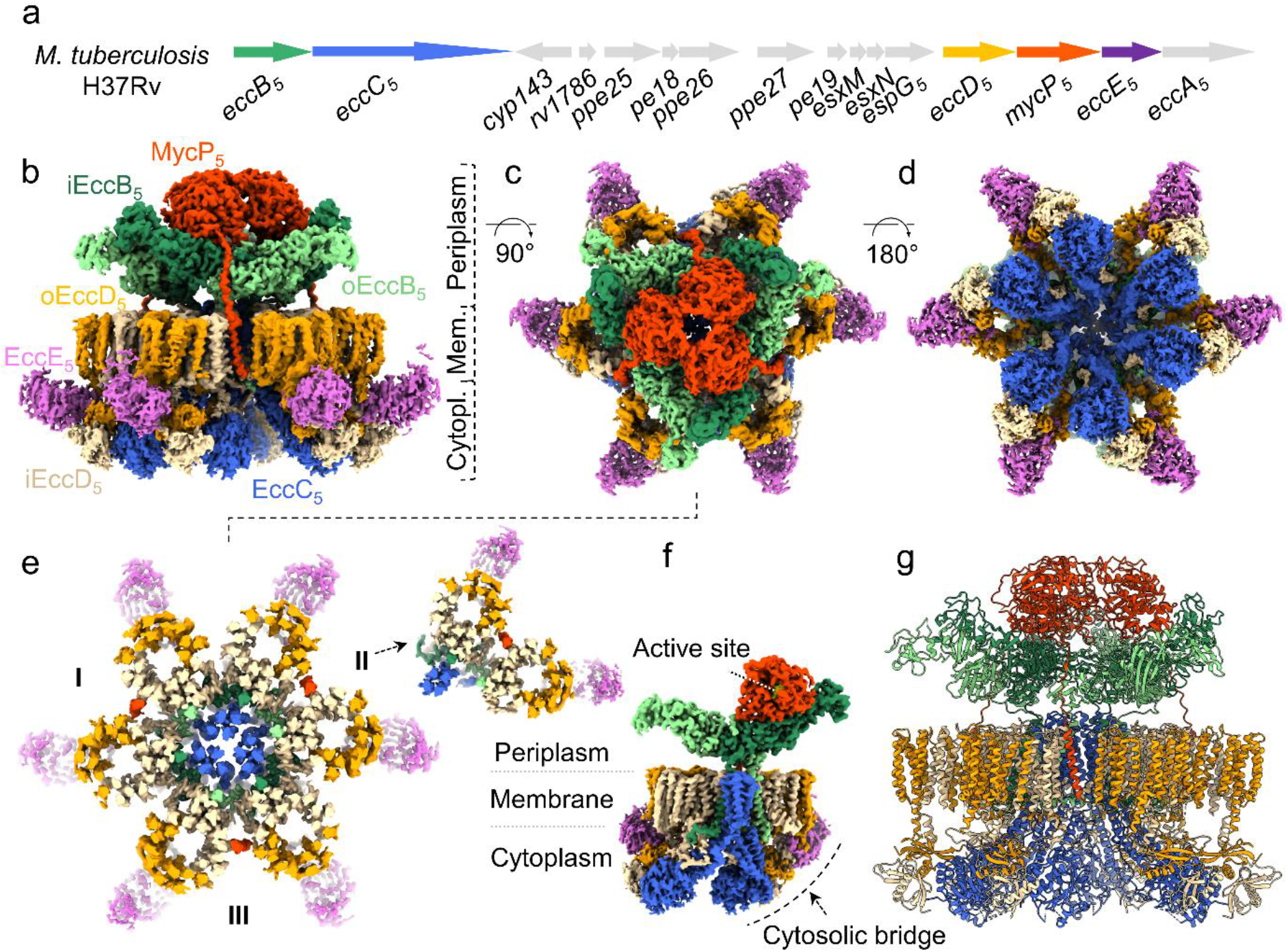
Cryo-EM structure of the intact ESX-5_mtb_ inner membrane complex. **a**, Genetic organization of the *esx-5* locus of *M. tuberculosis* H37Rv, which has been cloned and expressed in *M. smegmatis* MC^2^155. **b-e**, Cryo-EM density of the intact ESX-5_mtb_ inner membrane complex, zoned and coloured for every individual component. Components are: iEccB_5_, dark green; oEccB_5_, light green; EccC_5_, blue; iEccD_5_, beige; oEccD_5_, orange; EccE3, purple; MycP_5_, red. The full complex is 28.5 nm wide and 20 nm tall and has an absolute stoichiometry of 6:6:12:6:3 for EccB_5_:EccC_5_:EccD_5_:EccE3:MycP_5_. Side view (b), top view (c), bottom view (d) and a top cross section of the complex at membrane level, highlighting the 165 TMH region arrangement. For EccE_5_ only the cytoplasmic domain is shown, as the TMHs of this protein were poorly resolved. Inset contains a top cross section of an extracted dimeric unit, highlighting the TMH arrangement towards the potential closed pore. **f**, A single dimeric unit, viewed from the centre of the intact complex, highlighting the central four TMH-bundle formed by EccC_5_ and the position of MycP_5_ with its active site directed towards the inside of the periplasmic cavity. **g**, Ribbon model of the ESX-5_mtb_ assembly.

### Structural rearrangements of periplasmic domains

The periplasmic assembly of the ESX-5_mtb_ membrane complex is formed by six EccB_5_ copies organized in dimers and three MycP_5_ proteases. The three EccB_5_ dimers assemble in a triangle-shape, forming a central cavity (Fig. 1b; Fig. 2c). Two slightly different conformations of EccB_5_ can be distinguished, *i.e.* inner and outer EccB_5_ (iEccB_5_ and oEccB_5_, respectively), depending on their position. EccB_5_ dimerization is mainly mediated through its R1 and R4 repeat domains and further stabilization of the dimer is achieved by the EccB_5_ C-terminus, which wraps itself around its interacting EccB_5_ partner to form intermolecular hydrophobic contacts with its R1 domain (Fig. 2d). Central to these interactions is the 142-147 GIPGAP motif, a highly conserved region in all EccB homologs (Fig. 2d; Extended Data Fig. 15). Compared to the EccB3 dimer seen in the previously published ESX-3 subassembly^5,6^, the three EccB_5_ dimers are rotated by ~52 degrees with respect to their corresponding EccC_5_:EccD_5_:EccE_5_ membrane dimers (Extended Data Fig. 9d). This indicates that large conformational rearrangements are required during maturation into the fully assembled hexamer. To form the triangle-shaped assembly, iEccB_5_ engages oEccB_5_ of the adjacent dimer by packing its R3 domain against alpha helices α5 and α8 of domains R2 and R3, respectively, resulting in an asymmetric EccB_5_-tip arrangement (Fig. 2c). Due to the nature of this interaction, domain R3 of oEccB_5_ does not form any interactions at its tip extremity and thus displays higher flexibility, in line with previously observed crystallographic movements of this domain of monomeric EccB1 of *M. tuberculosis*^8^.

**Fig. 2.**
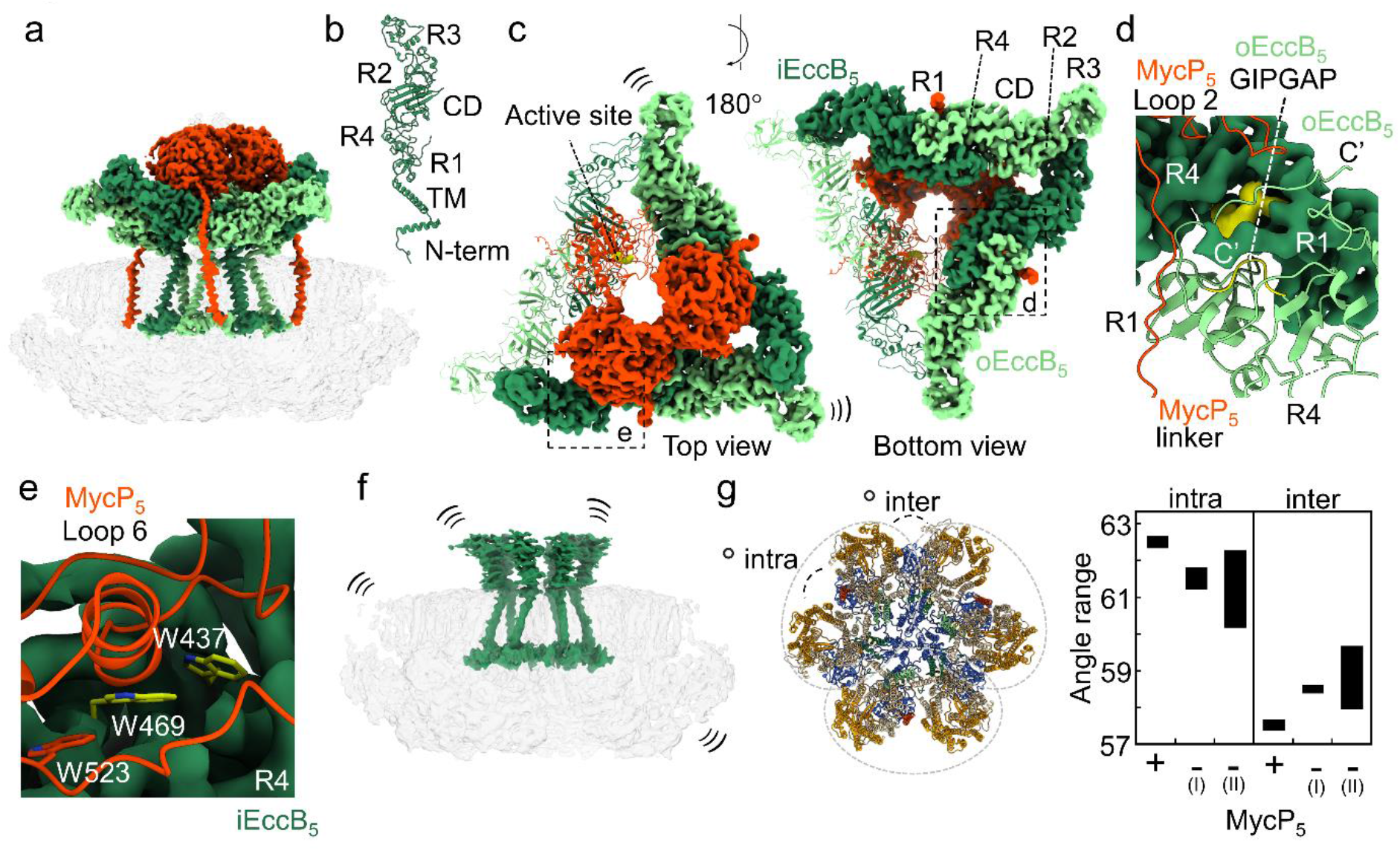
MycP_5_ acts as an allosteric driver for EccB_5_ hexamerization and membrane complex stabilization. **a**, Intact ESX-5_mtb_ assembly with EccB_5_ and MycP_5_ coloured as in Fig. 1 and other components transparent. **b**, Complete structure of monomeric EccB_5_mtb highlighting its overall fold and domains. **c** Top and bottom view of the EccB_5_-MycP_5_ periplasmic assembly with one unit (EccB_5_ dimer + MycP_5_ monomer) as ribbon model, highlighting the active site of MycP_5_ (in yellow) directed towards the inside of the periplasmic cavity. **d**, Zoom in on the EccB_5_ dimerization site highlighting the C-terminus of oEccB_5_ that is wrapped around the R1/R4 domain of the adjacent iEccB_5_ monomer, the conserved GIPGAP motif (in yellow) of EccB_5_ and the interactions of the EccB_5_ dimer with loop 2 and the linker domain of MycP_5_. **e**, Zoom in on the EccB_5_:MycP_5_ interaction surface, highlighting the three buried trypthophans. **f**, Map of ESX-5_mtb_ without co-purified MycP_5_ with EccB_5_ highlighted in dark green and the other components transparent. Three small, curved lines indicate the observed high flexibility of EccB_5_ at the periplasmic side and the overall heterogeneity of the membrane complex in the absence of MycP_5_. **g**, Angle variation range between protomers of the MycP_5_-bound (+) and two unbound states (-(I) and - (II)). Angles were calculated between the centre of masses of each individual protomer in relation to the centre of mass of the entire map. Intra, between two protomers within a dimer, i.e. that interact via the EccC_5_ four TMH-bundle in the central basket; Inter, between two protomers of adjacent dimers. In the MycP_5_-bound map, the periplasmic domain of EccB_5_ and full MycP_5_ proteins were removed for an accurate angle comparison between states.

### Three MycP_5_ subunits stabilize three EccB_5_ dimers into a periplasmic dome

Our periplasmic ESX-5_mtb_ map shows that the three MycP_5_ proteases together form a dome-like structure capping the periplasmic central cavity (Fig. 2a, c). Interactions between EccB_5_ and MycP_5_ are mainly mediated by the MycP_5_ protease domain and a composite interface generated by the R4 domain and loop 6 (Thr424-Ser435) of iEccB_5_ (Fig. 2e). The MycP_5_:EccB_5_ interface covers a surface area of ~1230 Å^2^, leading to the burial of three conserved tryptophans (EccB_5_ Trp437, Trp469 and MycP_5_ Trp523) (Fig. 2e). In addition, loop 2 of MycP_5_ binds to the C-terminus of oEccB_5_, explaining why a previous deletion of this loop showed reduced ESX-5 secretion in *M. marinum*^9^ (Fig. 2d, Extended Data Fig. 9k). Intermolecular MycP_5_:MycP_5_ interactions are mainly mediated through loop 1 and the N-terminal extension, running across the top of MycP_5_ protomers, and loop 3, which contacts the neighboring protease domain from the side (Extended Data Fig. 10e, f). MycP_5_ loop 5 (Ala151-Val271), which undergoes proteolytic processing during ESX-5 maturation^10^, folds up along the interface of two protease domains towards the central pore formed by the three MycP_5_ copies (Extended Data Fig. 10b). Although our cryo-EM map did not allow to build a complete model of loop 5 likely due to its high flexibility, at higher map thresholds this loop appears to cap the central periplasmic pore (Extended Data Fig. 10b). Notably, as loop 5 is not present in all mycosins and dispensable for ESX-5 secretion^9^, a speculative role in gating remains to be identified. Finally, the active sites of all three MycP_5_ proteases face towards the central lumen of the EccB_5_-MycP_5_ cavity (Fig. 2c), implying that potential substrates of this protease are translocated through and processed within this periplasmic chamber.

The dimer interface between i- and oEccB_5_ is by far the largest in the periplasmic EccB_5_-MycP_5_ assembly, covering a surface area of ~2000 Å^2^ and providing a solvation-free energy gain of ΔG = −23 kcal/mol per dimer upon complex formation (Fig. 2d). In contrast, the three additional EccB_5_ interfaces that are formed between the EccB_5_ dimers each cover a buried surface area of ~600 Å^2^ with a cumulative energy gain of only ΔG = −18 kcal/mol upon trimerization. This could provide an explanation as to why dimeric ESX subcomplexes are more stable as compared to their fully-assembled counterparts^5,6^. It is also noteworthy that the intermolecular EccB_5_:MycP_5_ interactions (with a surface area of ~395 Å^2^ and ΔG = −0.1 kcal/mol) are even more modest, providing a rationale as to why an interaction between MycP_5_ and the membrane complex has been elusive so far.

### MycP_5_ acts as an allosteric driver for EccB_5_ hexamerization and membrane complex stabilization

To further investigate the impact of MycP_5_ on the assembly of the EccB_5_ scaffold and its contribution to ESX-5_mtb_ stability, we analysed MycP_5_-free ESX-5_mtb_ complexes (Fig. 2f, g). These assemblies were present in our protein preparations and contained the same EccB_5_:EccC_5_:EccD_5_:EccE_5_ stoichiometry as the fully-assembled complexes (Extended Data Fig. 11). Following 3D reconstruction, two MycP_5_-free ESX-5_mtb_ maps displayed average resolution estimates of ~4.5 and ~6.7 Å (Extended Data Fig. 5, 6). The differences were most notable on the periplasmic side, where the six EccB_5_ copies failed to form a stable triangular scaffold in the absence of MycP_5_, but instead showed high flexibility (Fig. 2f). This shows that MycP_5_ acts as an allosteric driver that enables trimerization of the EccB_5_ dimers in the periplasm. This result is highly interesting, since mycosins are subtilisin-like proteases without any additional domains, apart from a TMH, some small loops and an N-terminal extension that wraps around the protein^11,12^. A more structural role of mycosin was predicted^7^ but now we finally understand the essential role of mycosins in T7SS.

In contrast to the periplasmic domain, the cytosolic and membrane regions were found to be similar to the MycP_5_-containing particles (Extended Data Fig. 11). However, these particles displayed an increased heterogeneity that reverberated throughout the complex, resulting into a slight waving of the membrane region and an increased angle variation between individual protomers (Fig. 2g, Extended Data Fig. 11). The presence of MycP_5_ synergistically reinforces not only the interactions between the periplasmic and membrane regions of the assembly but also between the two protomers of a dimer. As such, in the periplasm, MycP_5_ interacts mainly with one protomer via iEccB_5_, while in the membrane the TMH anchor of MycP_5_ interacts with the adjacent protomer of the dimer via oEccD_5_ (Extended Data Fig. 9f, g, 10).

### Hexamerization-induced formation of a membrane sealed central 12 TMH-bundle of EccC_5_ beneath the periplasmic dome

At the membrane level, six 22 TMH-containing EccD_5_ dimer barrels together form a circular raft with an inner cavity (Extended Data Fig. 12). Within this raft, inner EccD_5_ (iEccD_5_) monomers are situated closer to the center, whereas outer EccD_5_ (oEccD_5_) monomers are facing towards the periphery of the membrane complex. The EccD_5_ membrane barrels are structurally highly similar to the homologous EccD3 barrel in the ESX-3 subassembly^5,6^. The inner surface of each EccD_5_ barrel is decorated with densities attributable to stably-bound lipids or detergent molecules, suggesting that the EccD_5_ barrels in their native membrane environment are filled with membrane lipids (Extended Data Fig. 12e, f).

The single TMH of each EccB_5_ copy is anchored within the confinement of the EccD_5_ raft via hydrophobic interfaces provided by TMH6 and 11 of iEccD_5_ and stably-bound lipids (Fig. 3a, Extended data Fig. 9j). Notably, no substantial intermolecular interactions can be found between adjacent EccD_5_ barrels. Instead, coupling between two neighboring EccD_5_ barrels is achieved by the N-terminal loop and alpha helix of EccB_5_ that runs parallel to the cytoplasmic side of the IM and engages into interactions with the TMHs of the neighboring EccD_5_ barrel in a clockwise manner (Fig. 3a; Extended Data Fig. 9h, i). Because EccB_5_ TMH is slightly angled towards the center of the complex the architecture of the EccB_5_ hexamer at the membrane level is reminiscent of a basket, whose inner diameter shrinks from ~60 Å to ~45 Å when moving towards the periplasmic side (Fig. 3b).

**Fig. 3.**
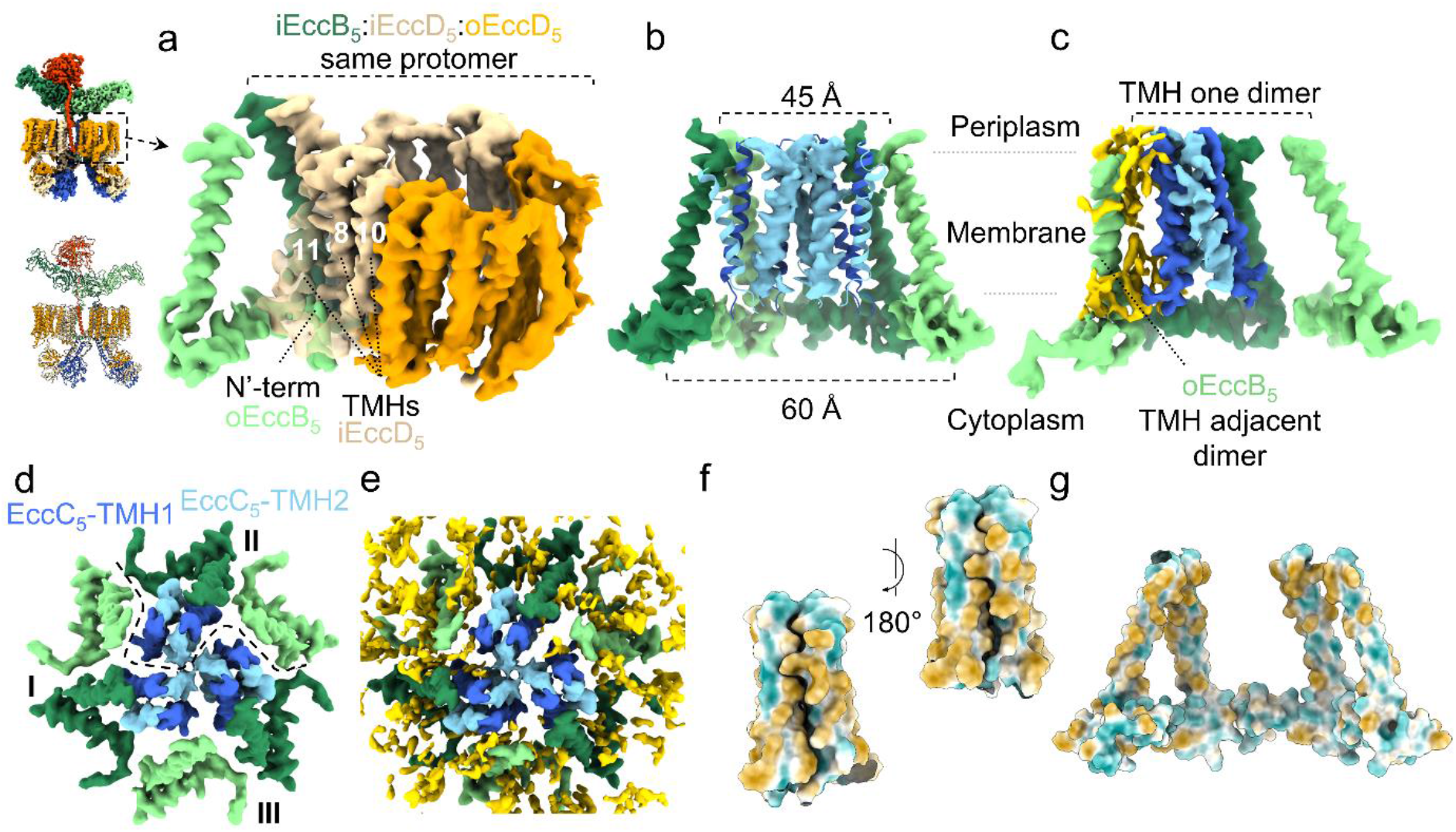
A basket formed by EccB_5_ TMHs holds three four TMH-bundles of EccC_5_. **a**, Angled view from the outside of the complex of the TMH and N-terminus of an oEccB_5_ interacting with a pocket formed by TMH 8, 10 and 11 of iEccD_5_ from the adjacent barrel. **b**, Side cross section through the EccB_5_ basket containing the three four TMH-bundles of EccC_5_. Light blue densities depict the three EccC_5_ TMH2s, one from each dimer, forming the central EccC_5_ TMH pyramid. Two EccC_5_ TMHs were removed for clarity. Sizes indicate the inner basket diameter at the cytosolic and the periplasmic side of the inner membrane. **c**, Side cross section through an EccB_5_ basket, showing that the EccC_5_ TMH-bundle does not interact with oEccB_5_ from its own dimer but forms lipid-mediated interactions with the oEccB_5_ TMH of the adjacent dimer. Lipids are shown in gold. **d**, Top view of the central EccB_5_ basket and the four EccC_5_ TMH-bundles. Dashed line marks the TMHs belonging to one dimeric unit. **e**, Same view as in d, highlighting the lipid-rich environment. In the central area surrounding the EccC_5_ pyramid, lipids are not clearly distinguishable, suggesting they are more mobile than lipids surrounding the EccB_5_ basket and EccD_5_ barrels. **f**,**g**, Surface model displaying the hydrophobicity of an EccC_5_ four TMH-bundle (f) and an EccB_5_ basket (g). Hydrophilic amino acids are shown in turquoise, hydrophobic residues in sepia.

The EccC protein is the only component present in all T7SSs, including related systems in Firmicutes^1^, and is therefore thought to be the central component in this nanomachinery. Each EccC protein has, in addition to 2 TMHs, four FtsK/SpoIIIE-like ATPase domains (also called nucleotide binding domains or NBDs) known to be important for secretion^5,6,13^. The twelve EccC_5_ TMHs could be fully resolved in the intact ESX-5_mtb_ complex and form three four-TMH bundles, each belonging to EccC_5_ molecules of one dimer (Fig. 3b, c, d; Extended Data Fig. 13). These bundles are held together by hydrophobic interactions and effectively seal the central space of the membrane complex. The central bundle is enclosed by the EccB_5_ TMHs, referred to as the EccB_5_ basket (Fig. 3b). Two EccC_5_ TMHs from each bundles contact the TMH of iEccB_5_, leaving the TMH of oEccB_5_ unbound by EccC_5_ (Fig. 3c, d). A pyramid-shaped assembly at the very center of the complex is formed by one TMH of each bundle (TMH2 of one EccC_5_ subunit), which aligns with the periplasmic EccB_5_-MycP_5_ chamber (Extended Data Fig. 13d).

The chambers within the confinement of the EccB_5_ basket appear to be filled with lipids (Fig. 3e). The density for these lipids is more ambiguous as compared to the stably-bound lipids in and around the EccD_5_ barrels, suggesting that these regions are more fluid. Consistently, although our full map reconstructions were centered on the TMHs of EccC_5_, the local resolution gradually increases when moving from the center to the EccB_5_ basket, where the resolution is highest (Extended Data Fig. 13c). This indicates that the three EccC_5_ four-TMH bundles display more flexibility as compared to the rigid EccB_5_ basket, a property that could come in favor when gating a secretion pore. The entrance to this putative pore widens on the cytoplasmic side where the EccC_5_ stalk domains expand radially (Extended Data Fig. 8, 13a). Together, our data suggests that six EccD_5_ barrels provide a stable scaffold for the assembly of a secretion pore, which is confined by EccB_5_ TMHs and gated via three EccC_5_ TMH-bundles. Secretion through this IM complex would require rearrangement of the flexible central TMHs of EccC_5_. Towards the periplasm, such a proposed central pore would extend into the periplasmic chamber, formed by EccB_5_ and MycP_5_.

### The cytoplasmic EccC_5_ domains resemble an extended and contracted conformation

At the cytoplasmic side of the complex, EccC_5_ has a stalk helix connecting its second TMH to the first NBD, also called the DUF/ATPase domain. This ATPase domain is bound to the cytosolic domains of i- and oEccD_5_, which, in turn, are bound to EccE_5_ at the periphery, together forming a so-called cytosolic bridge (Fig. 1e). Domains within this cytoplasmic bridge were challenging to model, due to the observed decreased connectivity. As a result of this, EccE_5_ could not be modelled confidently.

The distal C-terminal part of EccC_5_, which is composed of a string of three NBDs (1, 2 and 3), adopts two main conformations, herein called the extended and contracted conformation (Fig. 4a). In the extended state, the C-terminal three NBDs of each EccC_5_ bend parallel to the membrane, aligning below the cytosolic domains of oEccD_5_ and EccE_5_ of the same protomer and extending beyond the diameter of the membrane embedded assembly. Although the map did not permit model building in this area, the density can confidently accommodate a homology model consisting of the three EccC_5_ NBD domains (Fig. 4a; Extended Data Fig. 14). Further classification of this state reveals EccC_5_ to be heterogenous beyond NBD1, suggesting this to be more stably bound to components of its protomer. Although only a relatively small number of particles were found to be in the contracted conformation, the stable core of the membrane complex could be solved with sub nm resolution. (Extended Data Fig. 5). In the contracted state, the six flexible arms of EccC_5_ extend from the interface between the DUF/ATPase domain of EccC_5_ and the cytosolic domain of oEccD_5_ (Fig. 4a, Extended Data Fig. 14). Three separate disk-like structures can be observed, gradually constricting from the top to the bottom. This density shows a consistent gap at the interface between NBD1 and NBD2. This would allow the previously postulated binding of substrates to linker 2 connecting these two ATPase domains, resulting in displacement of this linker, required for NBD1 activation^6,14^. The highly dynamic cytoplasmic domains of the machinery may provide the basis for substrate selection, recognition or transport across the membrane.

In summary, our work highlights the fully-assembled structure of the ESX-5 inner membrane complex of *M. tuberculosis*. The structure will serve as a starting platform for the identification of important interactions that, once perturbed by small molecules, would aid in combating the important human disease tuberculosis.

**Fig. 4.**
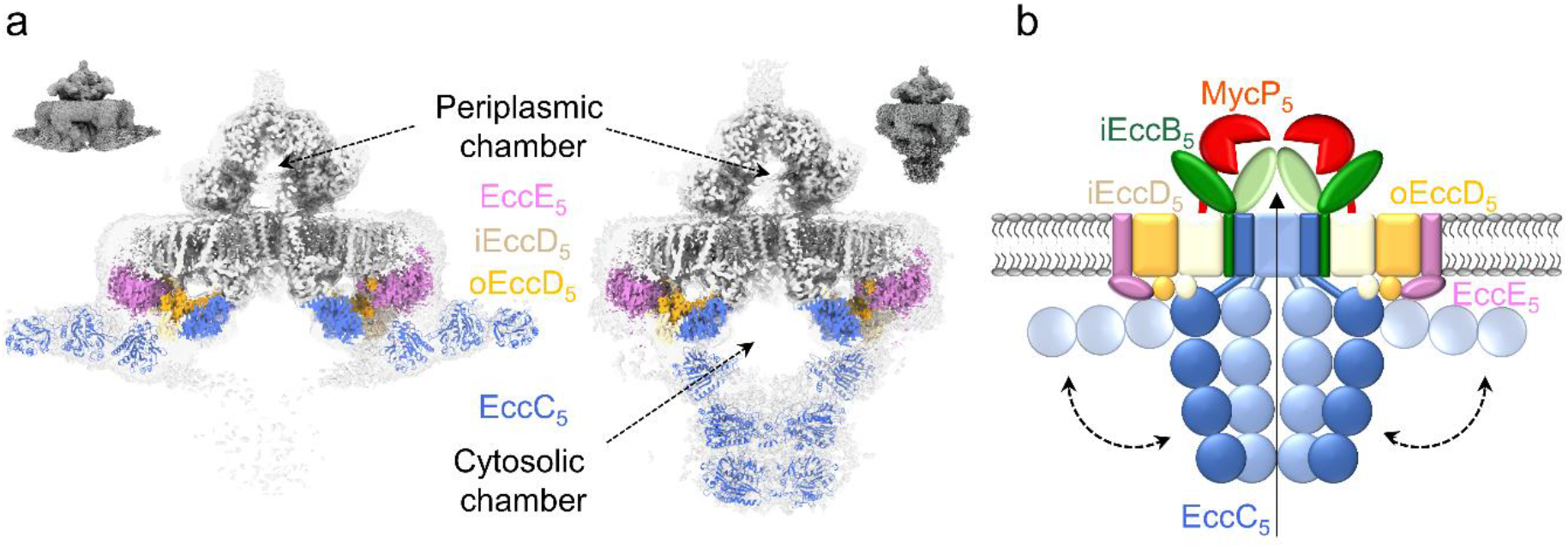
EccC_5_ adopts an extended and a contracted conformation. **a**, Side cross section of two density maps showing the extended (left) and contracted conformation (right) of EccC_5_. Highlighted are the periplasmic and cytoplasmic chambers formed by EccB_5_:MycP_5_ and by EccC_5_ upon closing. Homology models of the three C-terminal NBDs of EccC_5_ are fitted in the cytosolic densities. In the extended conformation, NBD1 of EccC_5_ localizes in close proximity to the cytosolic domains of iEccD_5_ and EccE_5_. Modelled EccC_5_ NBDs are shown in dark blue, the structures of the EccC_5_ DUF/ATPase domain are depicted in light blue and the EccE cytosolic domains are in purple. **b**, Model of the intact T7SS inner membrane complex, highlighting the two conformations of EccC_5_.

## Acknowledgements

We thank all members of the Marlovits, Houben and Bitter Laboratories for their support of this project. We would like to thank Wolfgang Lugmayr for scientific IT support and Luciano Ciccarelli for initial cryoEM analysis of the ESX-5_mtb_ complex. We thank Tristan Croll for his support with ISOLDE. High-performance computing was possible through access to the HPC at DESY/Hamburg (Germany). CryoEM data collection was performed at the CryoEM Facility at CSSB.

## Funding

This project was supported by funds available to TCM through the Behörde für Wissenschaft, Forschung und Gleichstellung of the city of Hamburg at the Institute of Structural and Systems Biology at the University Medical Center Hamburg-Eppendorf (UKE). The cryoEM facility is supported by the University of Hamburg, the University Medical Center Hamburg-Eppendorf, and DFG grant numbers (INST152/772-1 | 152/774-1 | 152/775-1 | 152/776-1 | 152/777-1 FUGG). This work received funding by a VIDI grant (864.12.006; to CMB and ENGH) from the Netherlands Organization of Scientific Research.

## Author contributions

CB, DF, JW, EH, WB, TCM designed experiments.

RU, CB generated constructs.

CB purified complexes; performed biochemical assays

JW vitrified samples and collected cryoEM images.

CB, JW collected negative stain images.

DF built the atomic model.

CB, DF, EH, WB, TCM interpreted data.

CB, TCM processed cryoEM data.

CB, DF, EH, WB, TCM wrote and revised the paper.

All authors read, corrected and approved the manuscript.

EH, WB, TCM supervised the project.

## Competing Interests

Authors declare no competing interests.

## Data and materials availability

Maps have been deposited at the EMDB database. Models have been deposited at the PDB.

## Material and Methods

### Molecular biology

*E. coli* Dh5α was grown at 37°C and 200 rpm in LB media supplemented with 30 μg/ml streptomycin. Cloning was performed in *E. coli* Dh5α using IProof DNA polymerase from BioRad and restriction enzymes from New English Biolabs. A list of the primers used for amplification can be found in Extended Data Table 3.

The plasmid expressing ESX-5_mtb_ was built as follows: the backbone of the previously described pMV ESX-5_mxen_ plasmid^1^ was modified to encode the unique restriction sites DraI and PacI upstream and SpeI and NdeI downstream of the TwinStrep tag sequence. The *rv1782-rv1783* (*EccB*_*5*_-*EccC*_*5*_) region, including ~380bp upstream of *EccB*_*5*_, of *M. tuberculosis* H37Rv was amplified while adding DraI and PacI restriction sites at the N and C-terminus, respectively (primers 1&2), and cloned into the modified plasmid upstream of the TwinStrep tag sequence, resulting in plasmid intermediate-1. The *M. tuberculosis* H37Rv region spanning *rv1791-rv1798* (*pe19-eccA5*) was amplified while adding SpeI and NdeI unique restriction sites (primers 3&4) and cloned downstream of the TwinStrep tag sequence into the intermediate plasmid, resulting in plasmid intermediate-2. Intermediate-2 plasmid was digested with SpeI and SnaBI, removing the region *rv1791-rv1794* (*pe19-espG5)*, and the region encompassing *rv1785-rv1794* (*cyp143-espG5*) was amplified as two individual PCR products (primers 5&6 and 7&8). The restricted backbone and PCR products were InFusion (Takara Bio) ligated, resulting in the final pMV ESX-5_mtb_ containing the entire *rv1782-rv1798* (*EccB_5_-eccA5*) locus.

### Isolation of mycobacterial cell envelopes

*M. smegmatis* MC^2^155 expressing ESX-5_mtb_ was grown at 37°C and 90 rpm in LB media supplemented with 0.05% Tween 80 and 30 μg/ml streptomycin. Cultures were grown to an OD_600_ of ~1.5, spun down for 15 minutes at 12.000xg in a JLA-8.1000 rotor and subsequently washed in PBS. After culture harvesting, all subsequent steps were performed at 4°C. Washed cell pellets were resuspended in buffer A (50 mM Tris-HCl pH 8, 300 mM NaCl and 10% glycerol) at a concentration of ~50 OD/ml and lysed by passing two times through a high-pressure homogenizer (Stansted) using a pressure of 0.83 kbar. Unbroken cells were pelleted at 5000xg for 5 minutes and supernatants were transferred to ultracentrifugation tubes. Cell envelopes were separated from the soluble fraction by ultracentrifugation at 150.000xg for 1.5 hours. Following ultracentrifugation, supernatants were discarded, pellets were washed once with buffer A, resuspended in buffer A at a concentration of 750-1000 OD/ml, snap-frozen in liquid nitrogen and stored at −80 °C until further use. The protein concentration of the cell envelop fraction was measured by BCA Assay (Pierce).

### Purification of the ESX-5_mtb_ membrane complex

The ESX-5_mtb_ membrane complex was purified in general as previously described^1^.

### Blue native polyacrylamide gel electrophoresis (BN-PAGE)

Samples consisting of either solubilized membranes or purified membrane complexes were mixed with 5% G-250 sample additive (Invitrogen), to a final concentration of ~0.2%, and run on 3-12% NativePage Bis-Tris Protein Gels (Invitrogen) according to manufacturer specifications. Gels were either transferred to PVDF membranes and stained with appropriate antibodies or stained with Coomasie R-250, both according to manufacturer specifications.

### Negative stain EM

Carbon-coated copper grids were glow discharged for 30 seconds at 25 mA using a GloQube^®^ Plus Glow Discharge System (Electron Microscopy Sciences). Four microlitres of diluted sample was applied to the grids and incubated for 30 seconds. The sample was blotted off from the side and the grid was washed briefly with 4 μl of staining solution (2% uranyl acetate) and then stained with 4 μl of the staining solution for 30 seconds. The stain was blotted off from the side and grids were air-dried. Grids were imaged using a Thermo Fisher Scientific Talos L120C TEM with a 4K Ceta CEMOS camera.

### Cryo-EM sample preparation

For the main dataset, purified sample was applied to Quantifoil R2/2, 200 mesh, copper grids floated with an additional ~1.1 nm layer of amorphous carbon. 4 μl of sample was applied onto glow-discharged grids (30 seconds at 25 mA) and allowed to disperse for 60 seconds at 4°C and 100% humidity. Grids were blotted for 4-6 seconds with a blot force of −5 and plunge frozen in a liquid propane/ethane mixture, using a Thermo Fisher Scientific Vitrobot Mark V. For the initial Arctica dataset, all steps were similar, with the exception of the additional layer of amorphous carbon.

### Cryo-EM - data acquisition

The initial cryo-EM dataset was collected on a 200 kV FEI Talos Arctica electron microscope equipped with a Falcon III direct electron detector running in counting mode and using Thermo Fisher Scientific EPU software. A total of 853 movies were recorded with a nominal magnification of 150.000x, corresponding to a pixel size of 0.96 at the specimen level. Movies were recorded with a total dose of 40.28 electrons/A^2^, fractionated in 38 frames over a 40 second exposure time and with a nominal defocus range of 1-2.5 μm.

The two high-resolution datasets were recorded using Thermo Fisher Scientific EPU software on a 300 kV Titan Krios TEM, equipped with a Gatan K3 direct electron detector running in counting mode and a Gatan Bioquantum energy filter (Slit size 10eV). 7984 and 9389 movies were recorded in counting mode in the two separate sessions with a nominal magnification of 81.000x, corresponding to a pixel size of 1.1 Å at the specimen level. Movies were recorded with a total dose of 59.5 electrons/A^2^, fractionated in 50 frames over a 3 second exposure time and with a nominal defocus range of 1-3 μm.

### Cryo-EM - data processing

Single particle analysis was performed using Relion3.1^2^, unless stated otherwise. For the initial Arctica dataset, movies were motion corrected, dose weighted and CTF was estimated using CTFFIND4^3^. Automated particle picking was performed using Cryolo^4^ and the pretrained Janni model. Following particle extraction and several rounds of 2D classification to remove obvious artifacts, an initial *de novo* model was generated. The dataset was further cleaned using 3D classification and the best class was subsequently used for reference-based particle picking. Following 2D/3D classification and 3D refinement in C1, the map displayed an apparent 3-fold symmetry and was further refined in C3. This final map displayed an estimated 13.5 Å resolution.

In the first Krios dataset, movies were motion corrected, dose-weighted and CTF estimated using CTFFIND4. Automated particle picking was performed using Cryolo with the pretrained Janni model and a low threshold. Particles were extracted and binned 4x and several rounds of 2D classification were performed followed by 3D classification with the 30 Å filtered Arctica model as a template. The resulting particles were re-extracted without binning, CTF corrected and polished and refined in C1, giving a map with an estimated overall resolution of 4.5 Å. For the periplasmic map, the center of mass for that region was determined using Chimera and the particles were recentered, extracted, CTF corrected, polished and 3D refined. Following refinement, density accounting to the periplasm was subtracted using a soft mask and refined in C1. For the cytosolic region, the center of mass of a cytosolic bridge dimer was determined using Chimera and particles were recentered, reextracted, CTF corrected, polished and 3D refined. Following refinement, the density accounting for individual cytosolic dimers was subtracted, resulting in a three times larger particle stack. Cytosolic dimers were first refined with a mask encompassing both cytosolic bridges. Subsequently, these were focus refined using a soft mask around one of the cytosolic bridges. This map was refined using the default Relion value “--tau2fudge 2” and also “--tau2fudge 4”, which increased the overall connectivity of the lower cytosolic area. The final map for the cytosolic bridge showed an estimated resolution of 3.3 Å and was sharpened using either Relion Postprocessing or DeepEMhancer. DeepEMhancer further helped in improving the observed anisotropy, overall map connectivity and feature resolving^5^.

The second Krios dataset was processed similarly, with some exceptions. Following 3D classification of the binned data against the 4.5 Å Krios map filtered to 30 Å, the two maps with and without MycP_5_ were processed separately. The MycP_5_-unbound map displayed increased heterogeneity and following unbinned re-extraction and refinement the particles were 3D classified without alignment, resulting in two obvious classes of 4.5 Å and 6.7 Å resolution. In contrast, a similar 3D classification for the MycP_5_-bound map did not further classify into structurally distinct classes. Following unbinned re-extraction and refinement, the MycP_5_-bound map showed an overall resolution of 4 Å which was further improved to 3.8 Å after C3 refinement. Model free density modification in Phenix.Resolve_Cryo_EM^6^ further improved the resolution of the entire C1 map to 3.8 Å and of the C3 refined map to 3.56 Å. The C1 map was further processed as previously described to obtain the periplasmic map at an estimated 3.8 Å resolution. Following 3D classification without alignment and further refinement in C3, the estimated resolution of the final periplasmic map improved to 3.5 Å. A final composite map was built using the 3.3 Å map of the cytosolic bridge, the 3.5 Å resolution map of the periplasmic portion and the membrane region of the C3 refined full map. The two states of EccC_5_ were separated by performing a masked 3D classification at the location of NBD1 of EccC_5_ on the 4 Å C1 full map. These particle subsets were subsequently recentered on the cytosolic region, reextracted with a bigger box size and refined.

### Model building and refinement

Model building started by generating homology models for MycP_5_, EccB_5_, EccC_5_ and EccD_5_ with Phyre2^7^. For MycP_5_, PDB entry 4J94^8^ served as a structural template, whereas PDB entries 4KK7^9^, 4NH0^10^/6SGW^11^ and 6SGZ^11^ served as reference models for EccB_5_, EccC_5_ and EccD_5_, respectively. To obtain atomic models of the periplasmic part (MycP_5_:EccB_5_) of the ESX-5_mtb_ complex, homology models of MycP_5_ and EccB_5_ were rigid body-fitted into a focus-refined periplasmic ESX-5_mtb_ map (EMDB: tba) using the fit-in-map tool in ChimeraX (v1.0)^12^. Model building, extension and interactive refinement was performed with ISOLDE (v.1.0.1)^13^, a molecular dynamics-guided structure refinement tool within ChimeraX (v.1.0). The resulting coordinate file (PDB: tba) was further refined with Phenix.real_space_refine (v.1.18.2-3874)^14^ using reference model restraints, strict rotamer matching and disabled grid search. Model validation was carried out using the MolProbity web server^15^ and EMRinger^16^ within the Phenix software package. Models for the membrane-embedded region (MycP_5_:EccB_5_:EccC_5_:EccD_5_) (PDB: tba) and cytoplasmic bridge (cytosolic domains of EccC_5_:EccD_5_) (PDB: tba) were built in the same way, using a reconstruction of the full ESX-5_mtb_ complex (EMDB: tba) and a focus-refined map of the cytoplasmic domains (EMDB: tba), respectively. The model of the periplasmic and the model of membrane-embedded region were subsequently fused and refined against the full ESX-5_mtb_ complex map (PDB: tba; EMDB: tba). Hereafter, a high-resolution composite map (EMDB: tba) was generated by fusing the full ESX-5_mtb_ complex map (EMDB: tba), zoned around the periplasmic and plasma-membrane areas, with six copies of the focus-refined cytoplasmic domains (EMDB: tba). Finally, a composite model (PDB: tba) was assembled.

Modeling into MycP_5_-free maps was performed with ISOLDE using the composite ESX-5 model in which MycP_5_ and the periplasmic domain of EccB_5_ (residues 84-507) were deleted. Adaptive distance restraints as well as torsion restraints were applied to all atoms to restrain short-range conformational changes but allow for long-range conformational movements. ISOLDE simulations for dynamic fitting of the coordinate file into EMDB tba and EMDB tba were performed (~10 min, 10 Kelvin) after which the models showed satisfying fits to the new maps without further manual intervention.

Visualization of atomic coordinates and map volumes was performed with ChimeraX (v.1.1).

## Extended Data Figures

**Extended Data Fig. 1.**
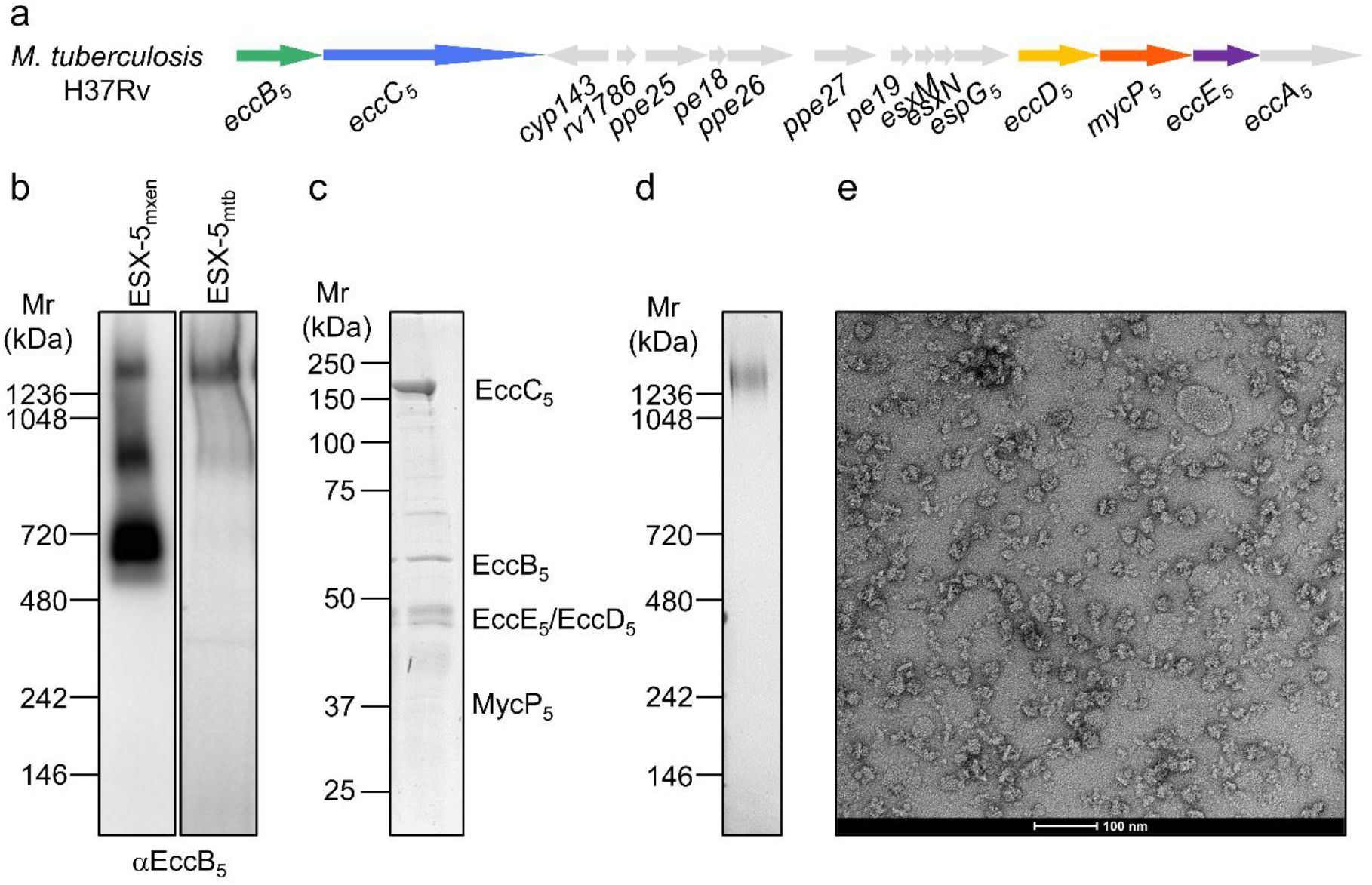
Initial purification of the ESX-5_mtb_ membrane complex. **a**, Genetic organization of the *esx-5* locus of *M. tuberculosis* H37Rv, which has been cloned and expressed in *M. smegmatis* MC^2^155. **b**, Blue native PAGE and western blot analysis using an anti-EccB_5_ antibody of DDM-solubilized membranes from *M. smegmatis* MC^2^155 expressing ESX-5_mxen_ and ESX-5_mtb_. **c**, **d**, SDS-PAGE (c) and BN-PAGE (d) and Coomassie staining of Strep- and SEC-purified ESX-5_mtb_ membrane complexes. **e**, Negative-stain EM analysis of ESX-5_mtb_ membrane complexes shown in c and d.

**Extended Data Fig. 2.**
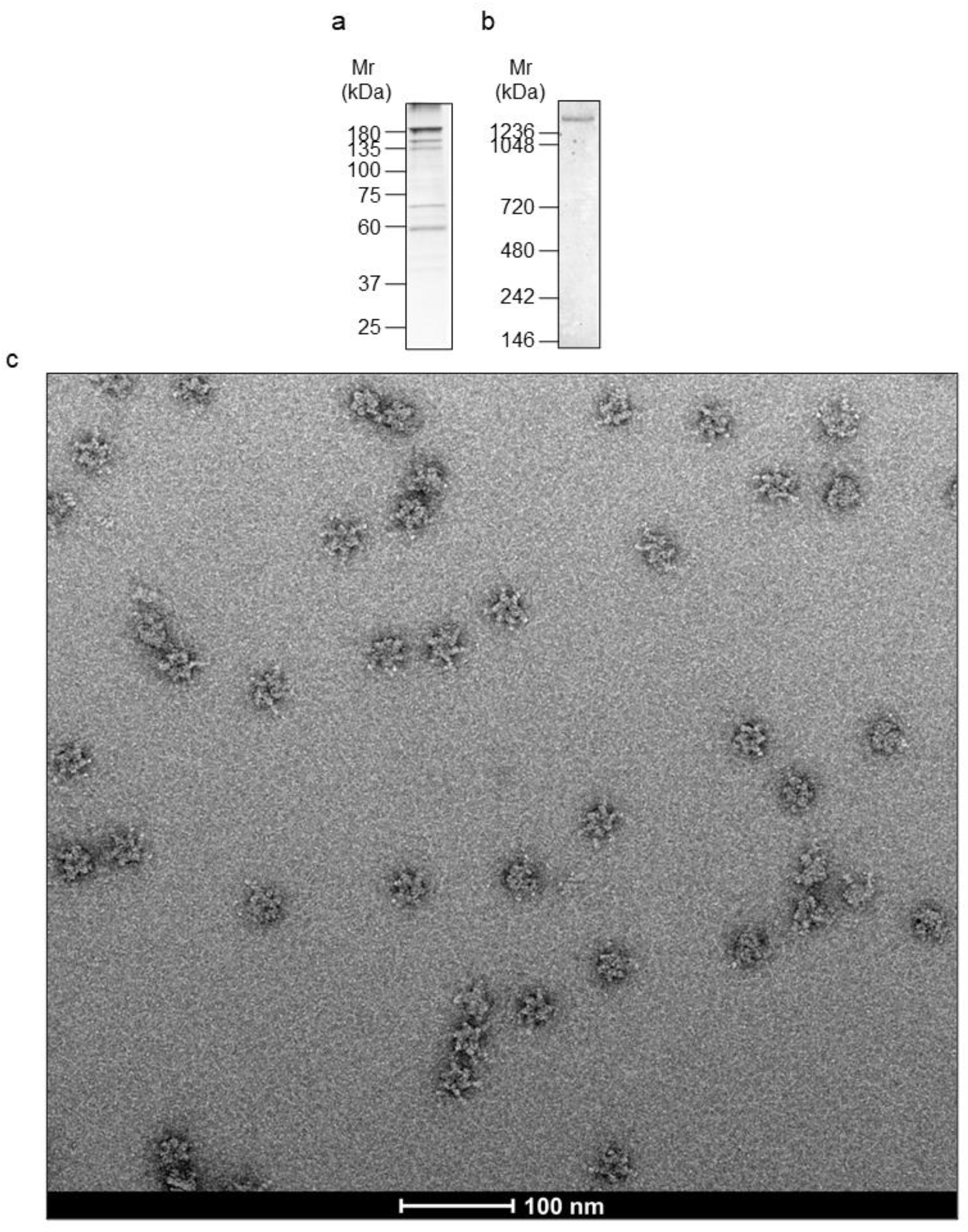
Final purification of the ESX-5_mtb_ membrane complex. **a**, **b**, SDS-PAGE (c) and BN-PAGE (d) and Coomassie staining of Strep- and SEC-purified ESX-5_mtb_ membrane complexes final sample. **c**, Negative-stain EM of the same sample as in a and b.

**Extended Data Fig. 3.**
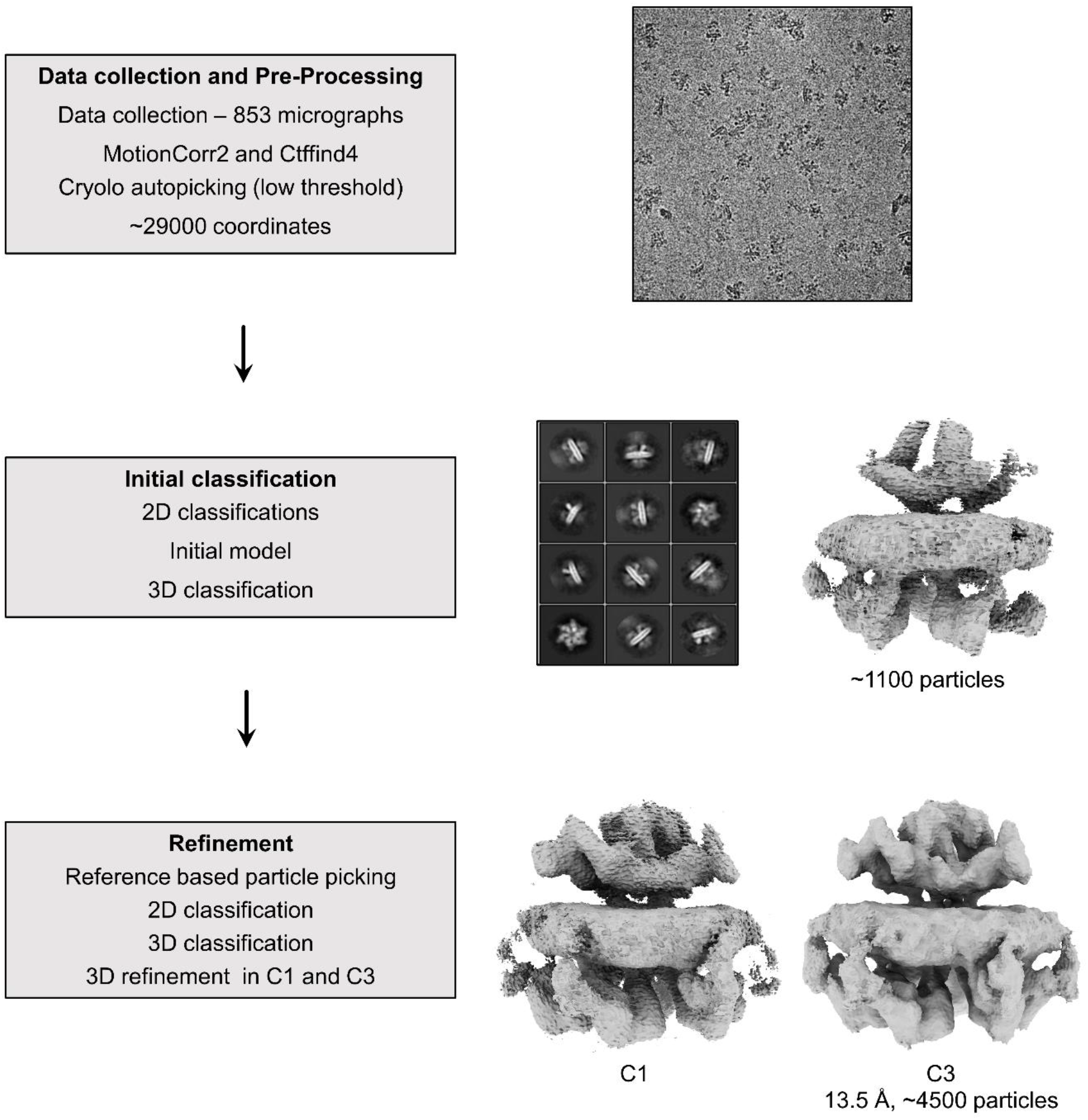
Cryo-EM data collection and single particle reconstruction procedure for the initial Talos Arctica collected dataset.

**Extended Data Fig. 4.**
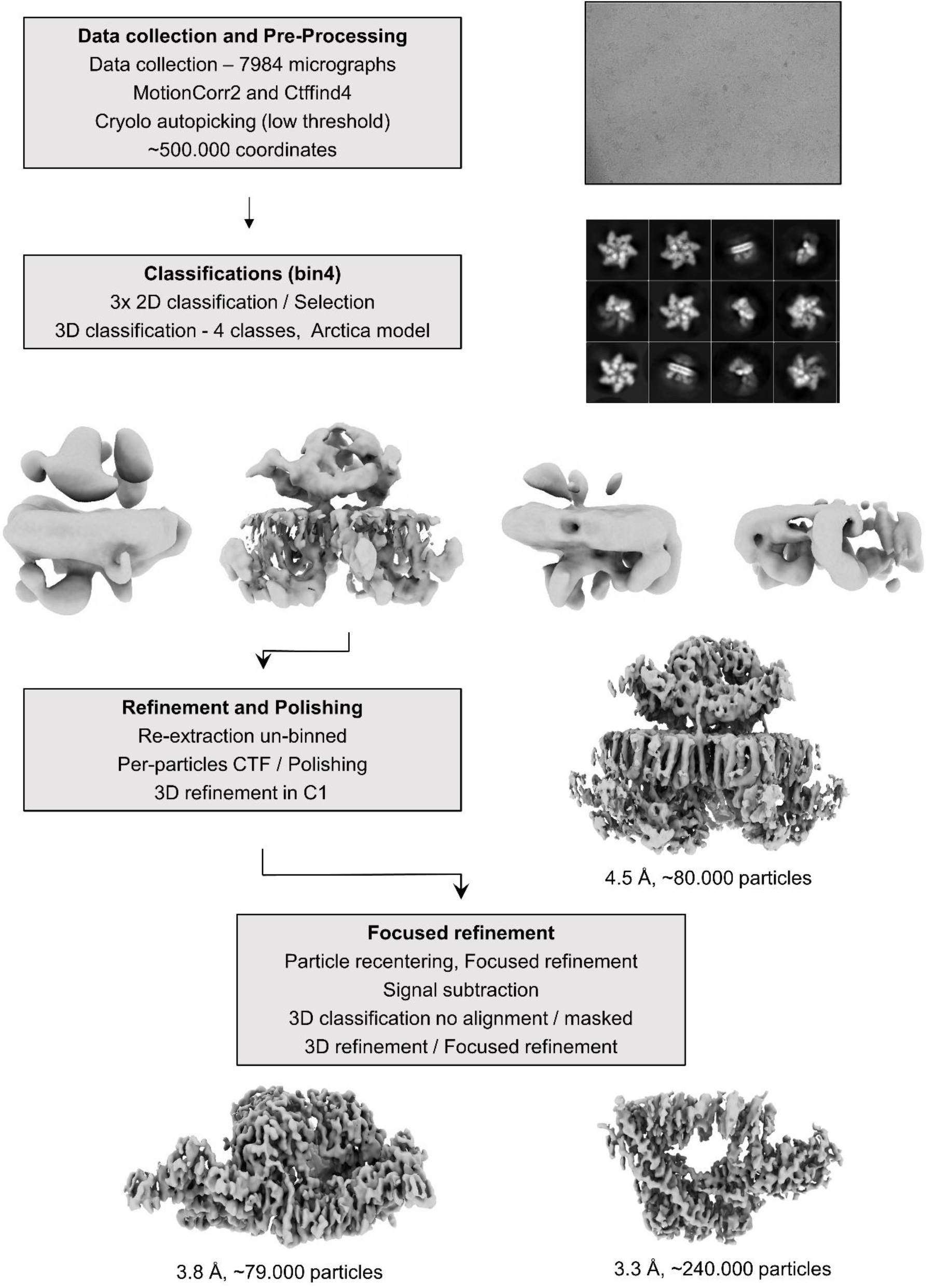
Cryo-EM data collection and single particle reconstruction procedure for the first high-resolution Titan Krios collected dataset.

**Extended Data Fig. 5.**
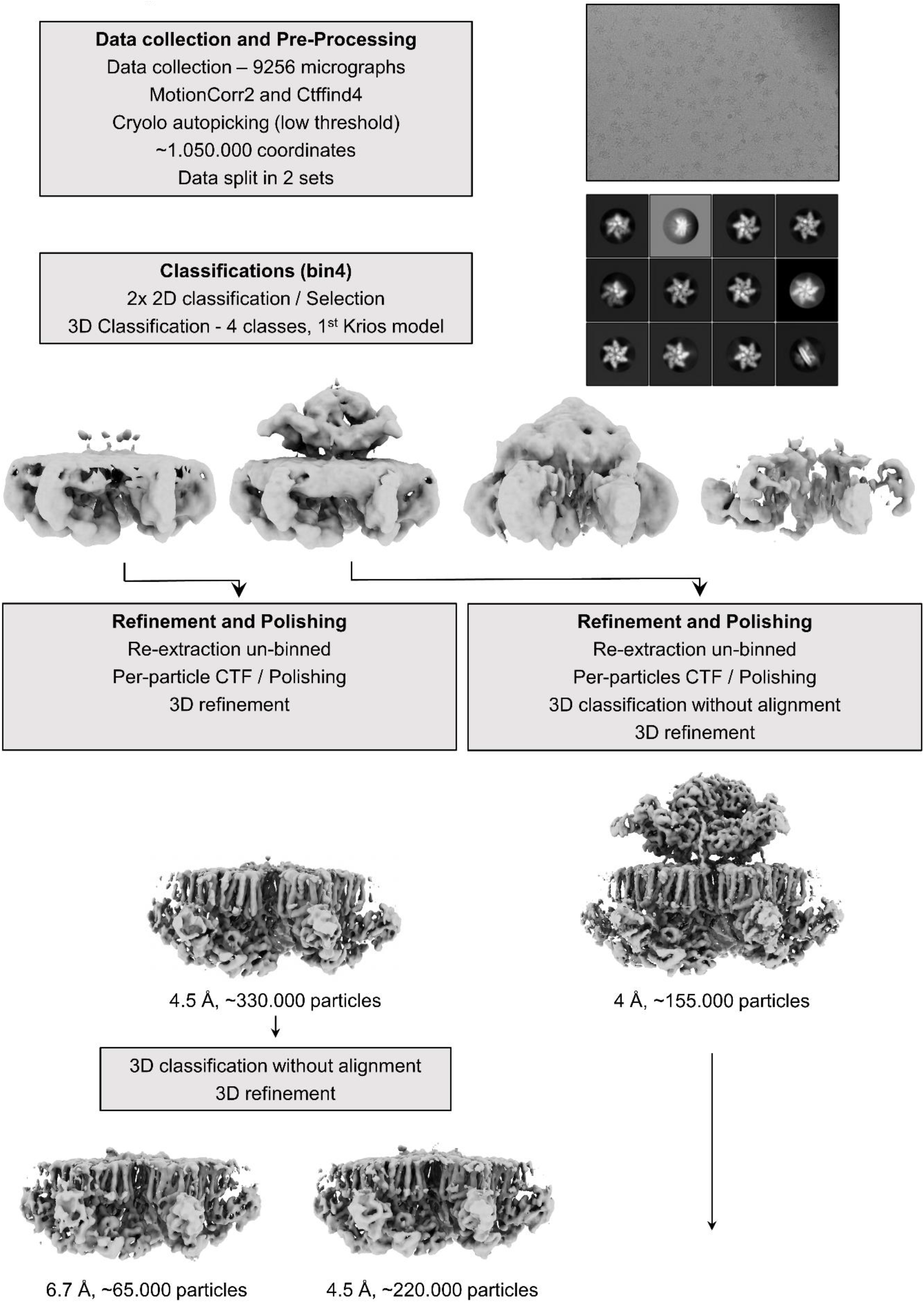
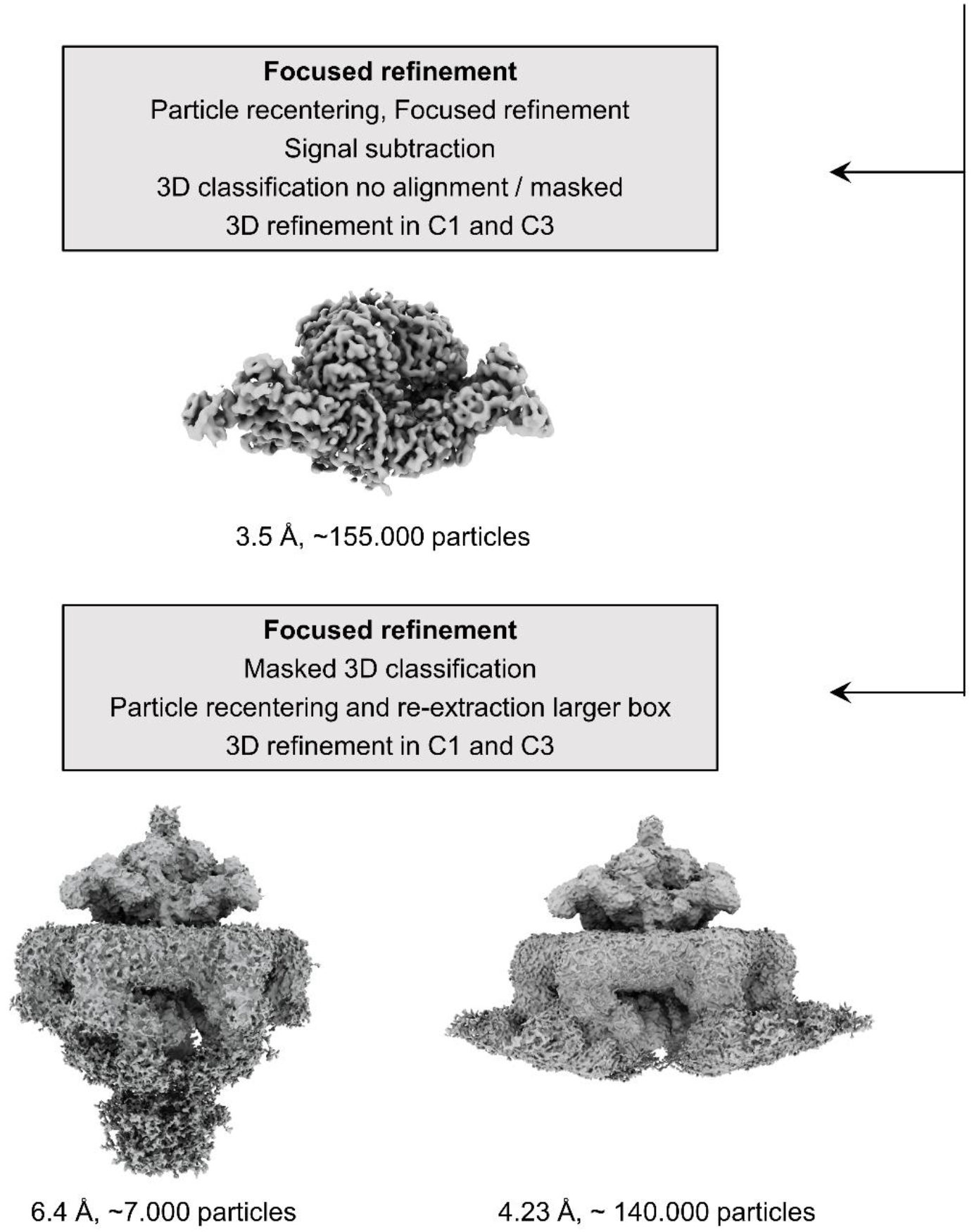
Cryo-EM data collection and single particle reconstruction procedure for the second high-resolution Titan Krios collected dataset.

**Extended Data Fig. 6.**
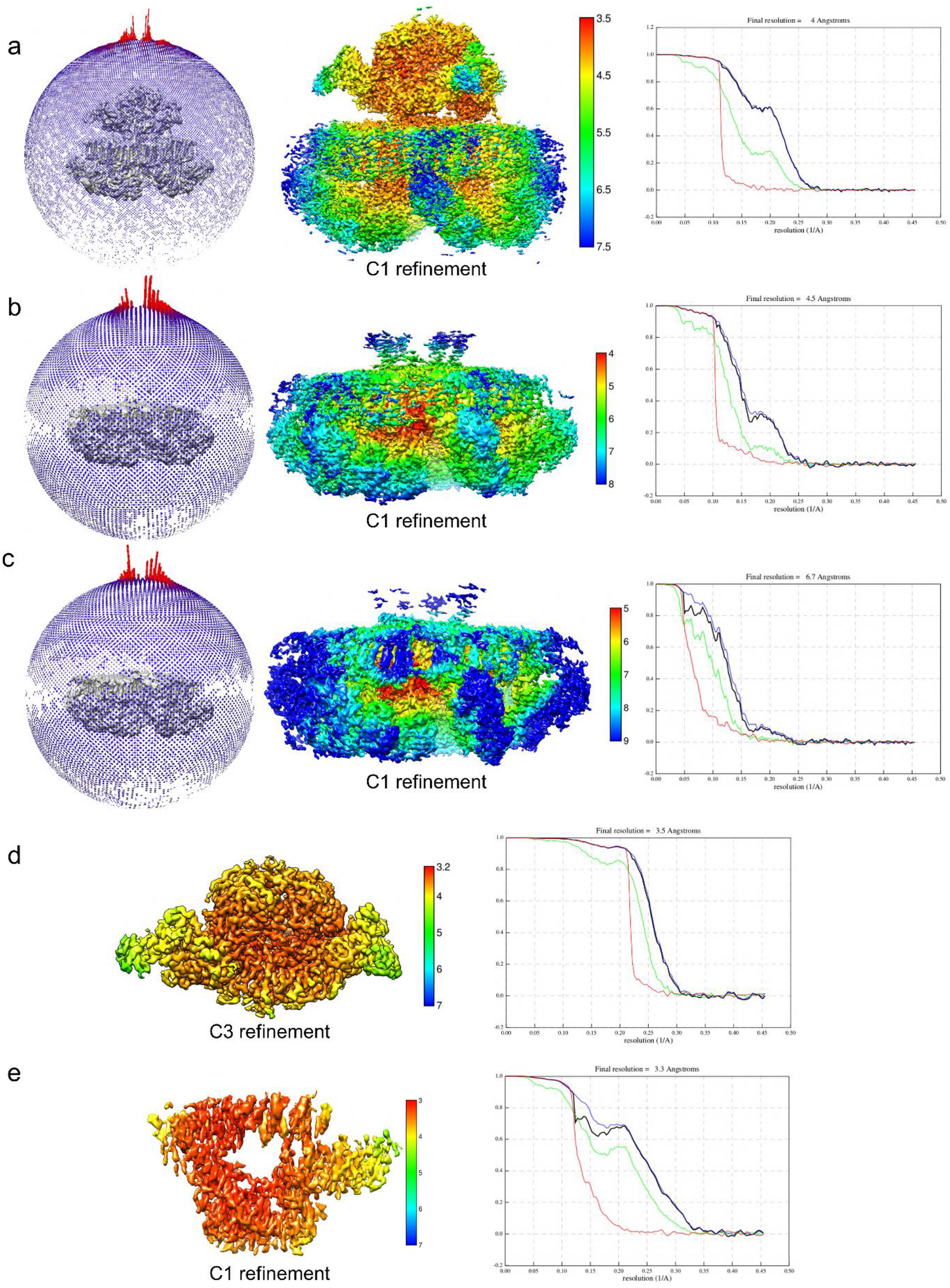
Single particle reconstructions of the ESX-5_mtb_ membrane complex. **a**-**c**, Angular distribution plots, local resolution estimations and Fourier Shell Correlation (FSC) plots of C1 reconstruction of the entire MycP_5_ bound ESX-5_mtb_ membrane complex (a), C1 reconstructions of the two heterogenous MycP_5_ unbound ESX-5_mtb_ membrane complexes (b, c). **d**, **e**, Local resolution estimation and FSC plot for the C3 refined periplasmic map (d) and the map of the cytosolic bridge (e).

**Extended Data Fig. 7.**
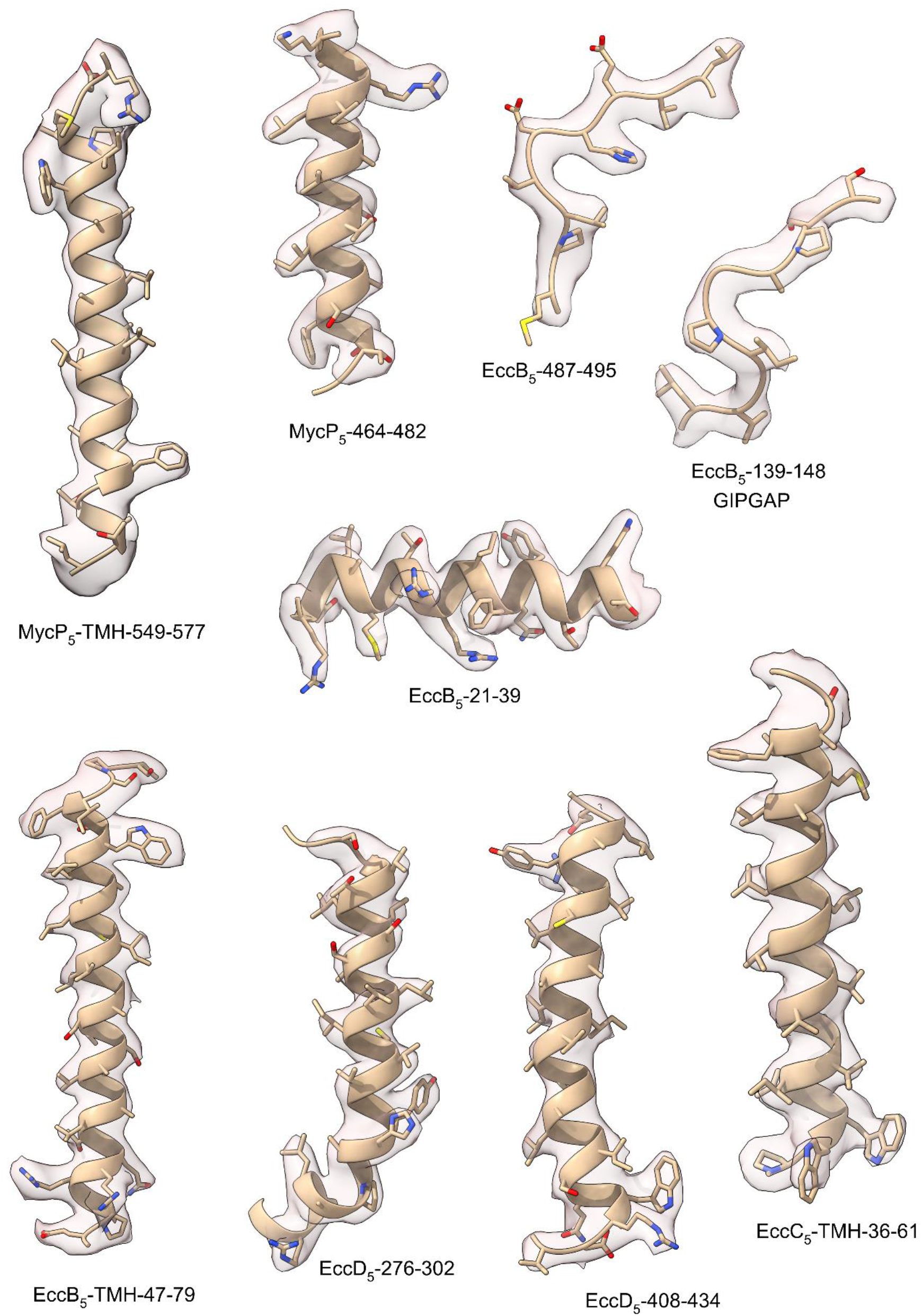
Examples of Cryo-EM densities.

**Extended Data Fig. 8.**
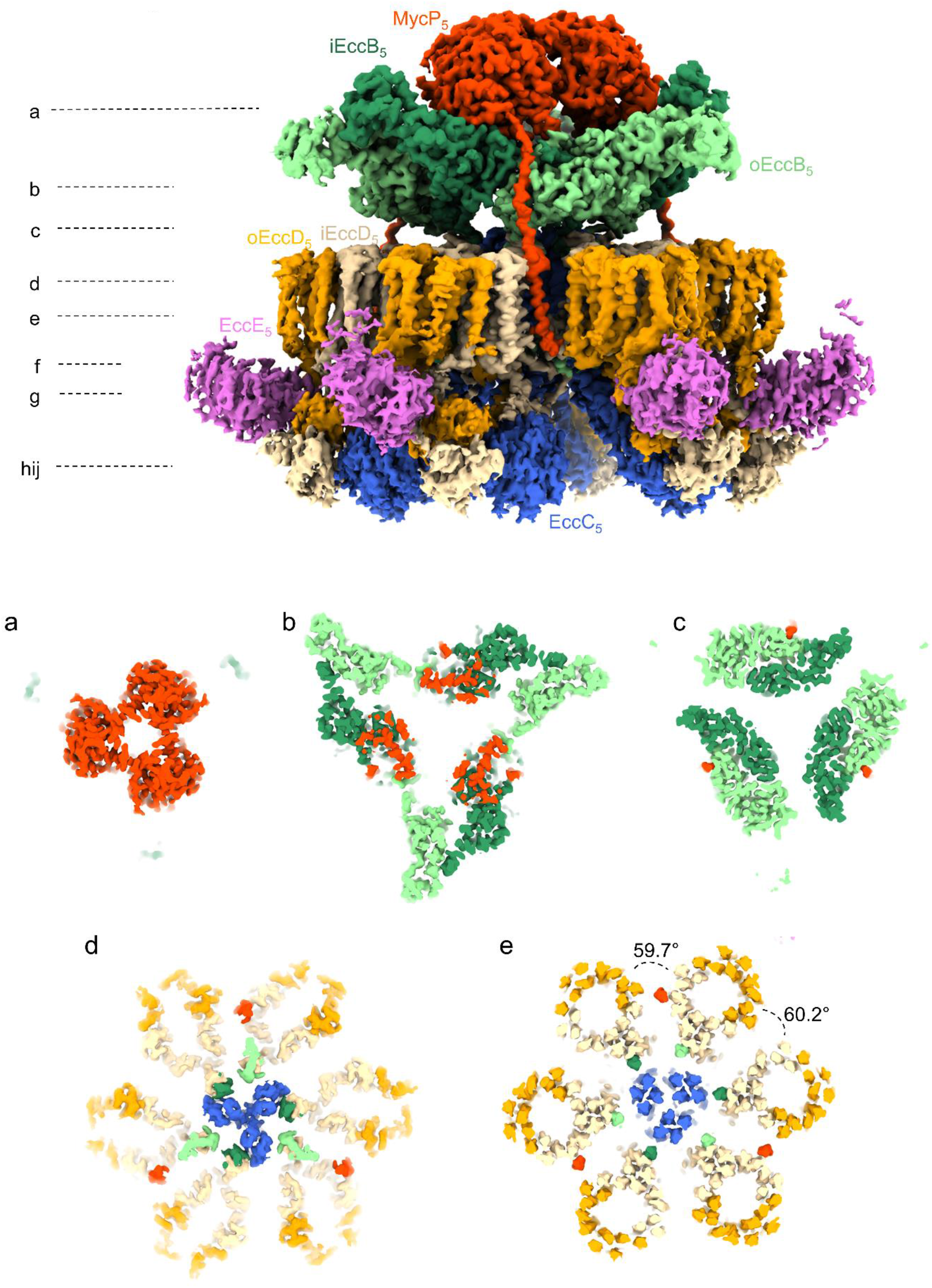
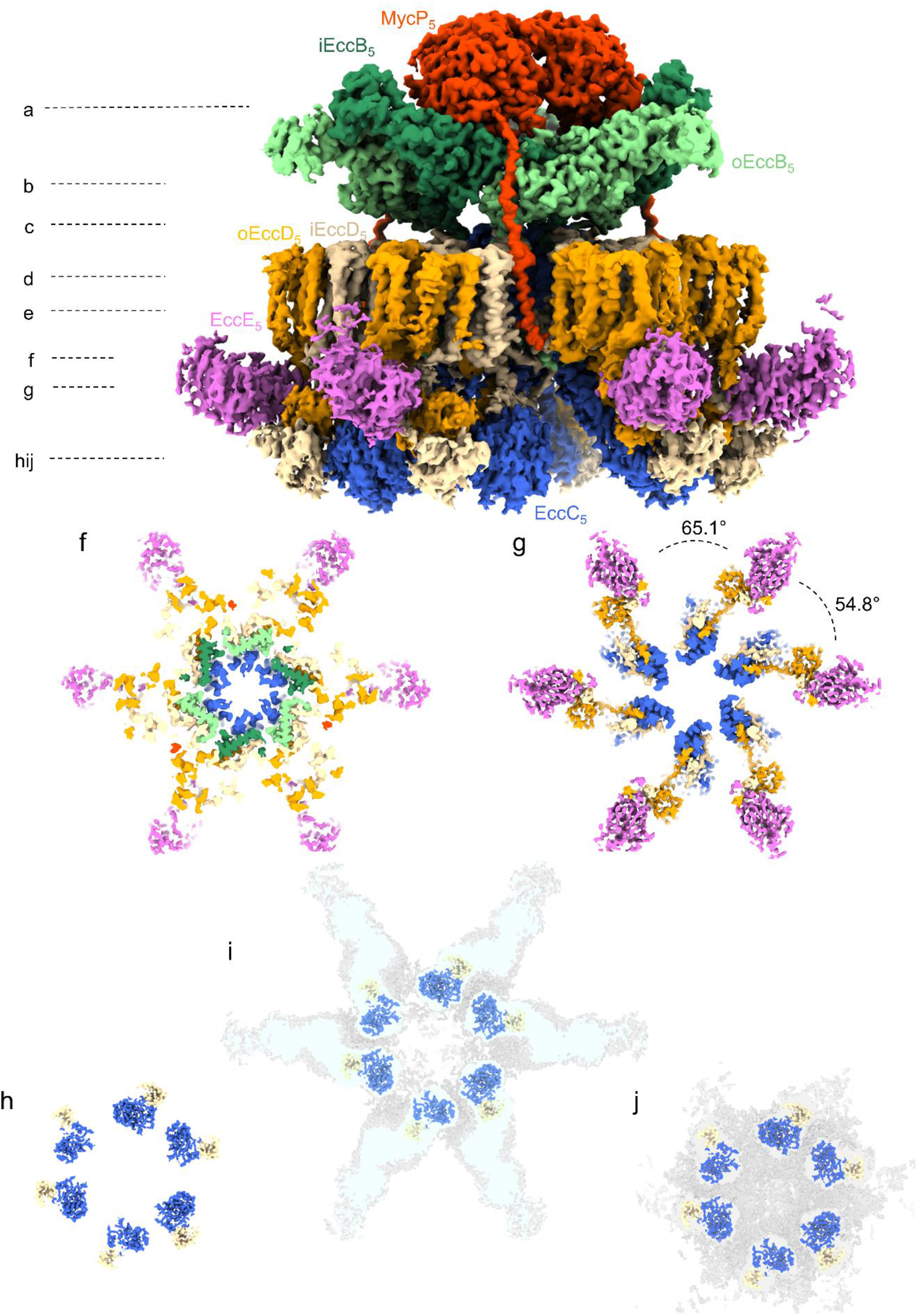
Top cross sections through the intact ESX-5_mtb_ membrane complex. **a**, MycP_5_ trimer top view, highlighting the pore formed at the periplasmic side. **b**, Section through the periplasmic assembly at the EccB_5_:MycP_5_ interface, showing the position of the protease domain sitting on top of iEccB_5_. **c**, Section through the periplasmic assembly at the EccB_5_ dimer interface level, highlighting the MycP_5_ linker connection to the TMH. **d**, Top view of the six membrane protomers with the closed EccC_5_ TMH-pyramid at the centre. **e**, Top cross section through the six membrane protomers highlighting 153 of the 165 TMHs. At the central area, towards the cytosol, the three EccC_5_ TMH-pyramid opens up like an iris. MycP_5_, whose protease domain interacts with the protomer containing iEccB_5_, interacts with the oEccD_5_ barrel of the adjacent protomer at the membrane level. At the membrane level, the angle between protomers within a dimer and between adjacent protomers of different dimers differs by only 0.5 degrees. **f**, Top section displaying the region below the inner leaflet of the inner membrane highlighting a further opening of the EccC_5_ gated pore and the lower part of the EccB_5_ basket, formed by EccB_5_ N-termini. **g**, At the cytosolic level, the angle between protomers differs to that at the membrane level. As such, the angle between protomers within a dimer grows with 5.4 degrees, while the angle between adjacent protomers of different dimers decreases by 5.4 degrees. **h**, Section through the lower region of the cytosolic bridge, containing the DUF/ATPase domain of EccC_5_ and the cytosolic domain of iEccD_5_. **i**, Same view as in h, but then overlayed with the EccC_5_ extended state, highlighting the radial extension of the EccC_5_ NBD1-3 almost parallel to the inner membrane. **j**, Same view as in h, but then overlayed with the EccC_5_ contracted stated.

**Extended Data Fig. 9.**
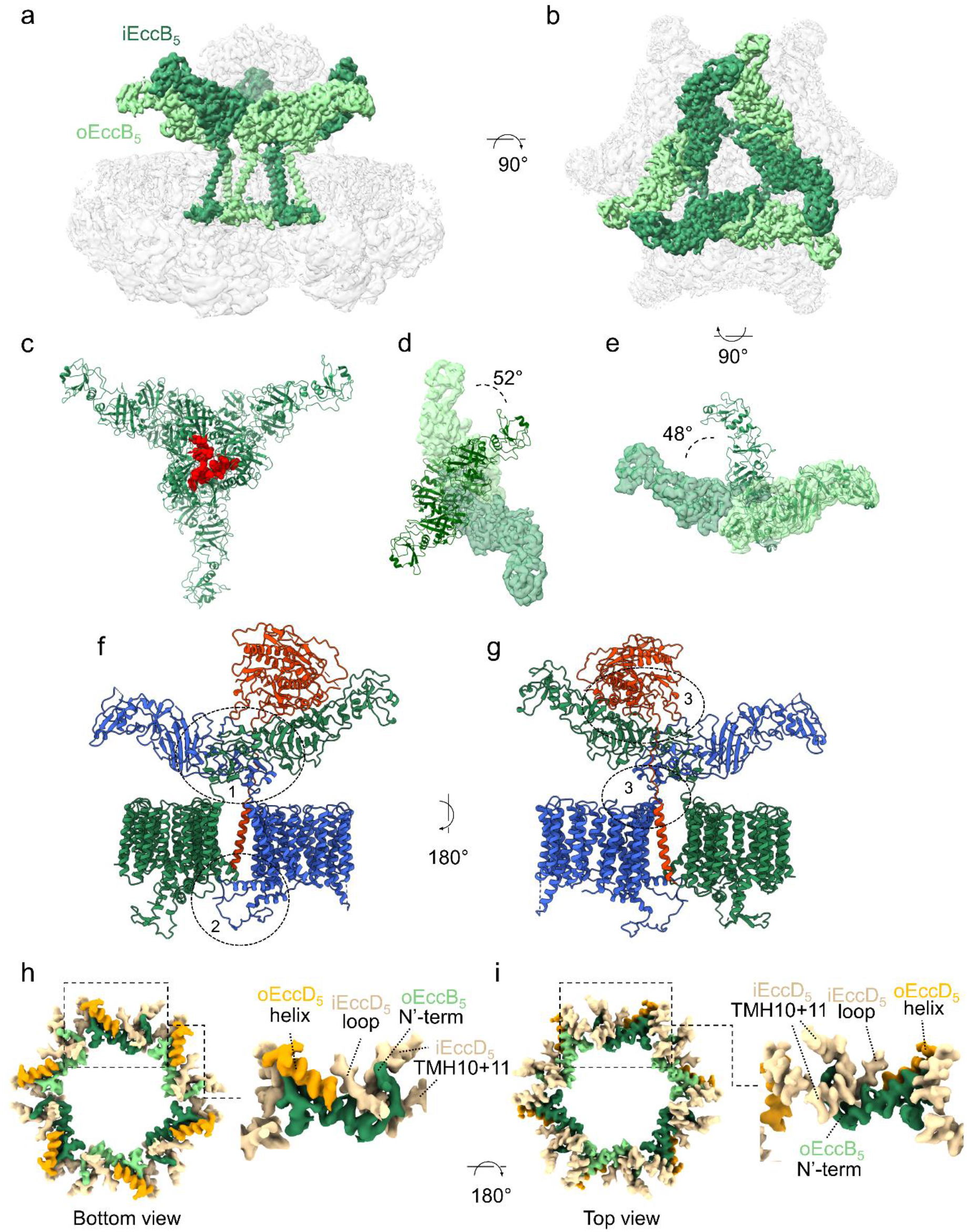
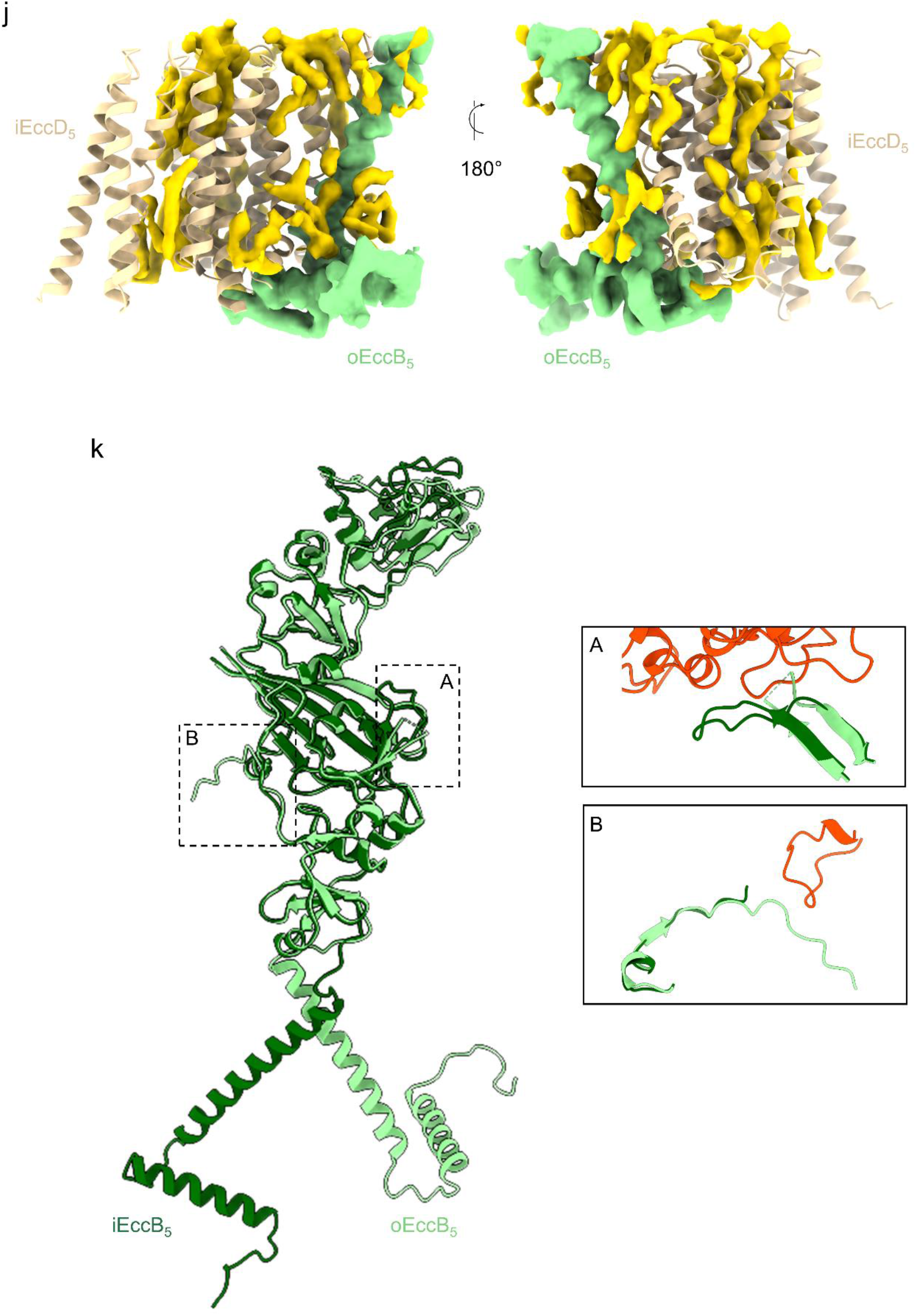
Hexameric EccB_5_ adopts a triangular conformation in the periplasm. **a**, **b**, Side (a) and top (b) view of an intact ESX-5_mtb_ assembly in which i- and oEccB_5_ are coloured as in Fig. 1 and the rest of the components are transparent. **c**, A V-shaped EccB3 dimer (PDB: 6sgy) was fitted into the *M. smegmatis* ESX-3 dimer cryo-EM density (EMD: 20820) together with the corresponding dimeric ESX-3 model for the membrane and cytosolic domains (PDB: 6UMM) using the Chimera Fit in Map tool. This combined model was subsequently trimerized, based on our full ESX-5 map reconstructions. The clashing of EccB3 periplasmic domains between the dimers towards the central area in this hybrid model are highlighted in red. **d**, Upon MycP_5_ binding to the entire assembly, the periplasmic EccB_5_ dimer is rotated with 52 degrees, avoiding the clashes observed in c. Angles were measured by aligning the hybrid model and the ESX-5_mtb_ model at the membrane level. Subsequently, centres of mass were defined for the combined R1 domains of each dimer (at the base of the dimer) and for every R2/R3 EccB monomer (towards the tips of the EccB dimer). Planes defined by these three points were generated for both EccB3 and EccB_5_ dimers and angles were measured between these two planes. **e**, Compared to the V-shaped EccB3 dimer, the angle between the two EccB_5_ monomers grows with 48 degrees upon MycP_5_ binding. EccB dimer angles were calculated by measuring the angle between the centres of mass of the R2/R3 domains of each EccB_5_ protomer in relation to the centre of mass of both R1 domains. **f**, **g**, Inside (f) and outside (g) view of a dimer containing EccB_5_, MycP_5_ and the TMHs of EccD_5_. For clarity purposes, one EccD_5_:EccB_5_ protomer is coloured in blue, the second protomer in green and MycP_5_ in red. At dimer level, three cross protomer contact points can be observed upon MycP_5_ binding: one at the periplasmic level, one at the cytosolic level and one between the periplasmic region and the TMHs. At the periplasmic level, the two EccB_5_ monomers interact via their R1/R4 domains and C-termini that wrap around their respective EccB_5_ partners. At the cytosolic level, the N-terminus of EccB_5_ interacts with a pocket formed by TMH 8, 10 and 11 of iEccD_5_ of the adjacent protomer. The membrane to periplasm cross protomer interaction is realized through MycP_5_. At the periplasmic level, MycP_5_ mainly interacts through, its protease domain, with one protomer via iEccB_5_. In addition, the TMH of MycP_5_ interacts with iEccD_5_ of the adjacent protomer, towards the periplasmic leaflet of the inner membrane. **h**, Bottom view of the lower cytosolic area of the EccB_5_ basket, formed by EccB_5_ N-termini (10-48) and depicted with the interacting pocket formed by TMH 10, 11 (456-477) and 8 (not shown for clarity) of iEccD_5_ of the adjacent protomer. The EccB_5_ N-terminus is also buttressed in this position by a short helix (119-130) of oEccD_5_, which connects the EccD_5_ barrel with its cytosolic domain, and also by part of the iEccD_5_ loop (307-315) that subsequently folds along the stalk and DUF/ATPase domain of EccC_5_. **i**, Same map as in h, but viewed from the top. **j**, Side views of the TMH region of iEccD_5_, depicted as a ribbon model, and the TMH and N-terminus of an interacting oEccB_5_, depicted as zoned density. An array of lipids found in the EccD_5_ barrel but also surrounding this iEccD_5_:EccB_5_ interaction site are depicted in gold. **k**, Superposition of iEccB_5_ and oEccB_5_, highlighting conformational differences between the two, which are the result of the interaction with MycP_5_.

**Extended Data Fig. 10.**
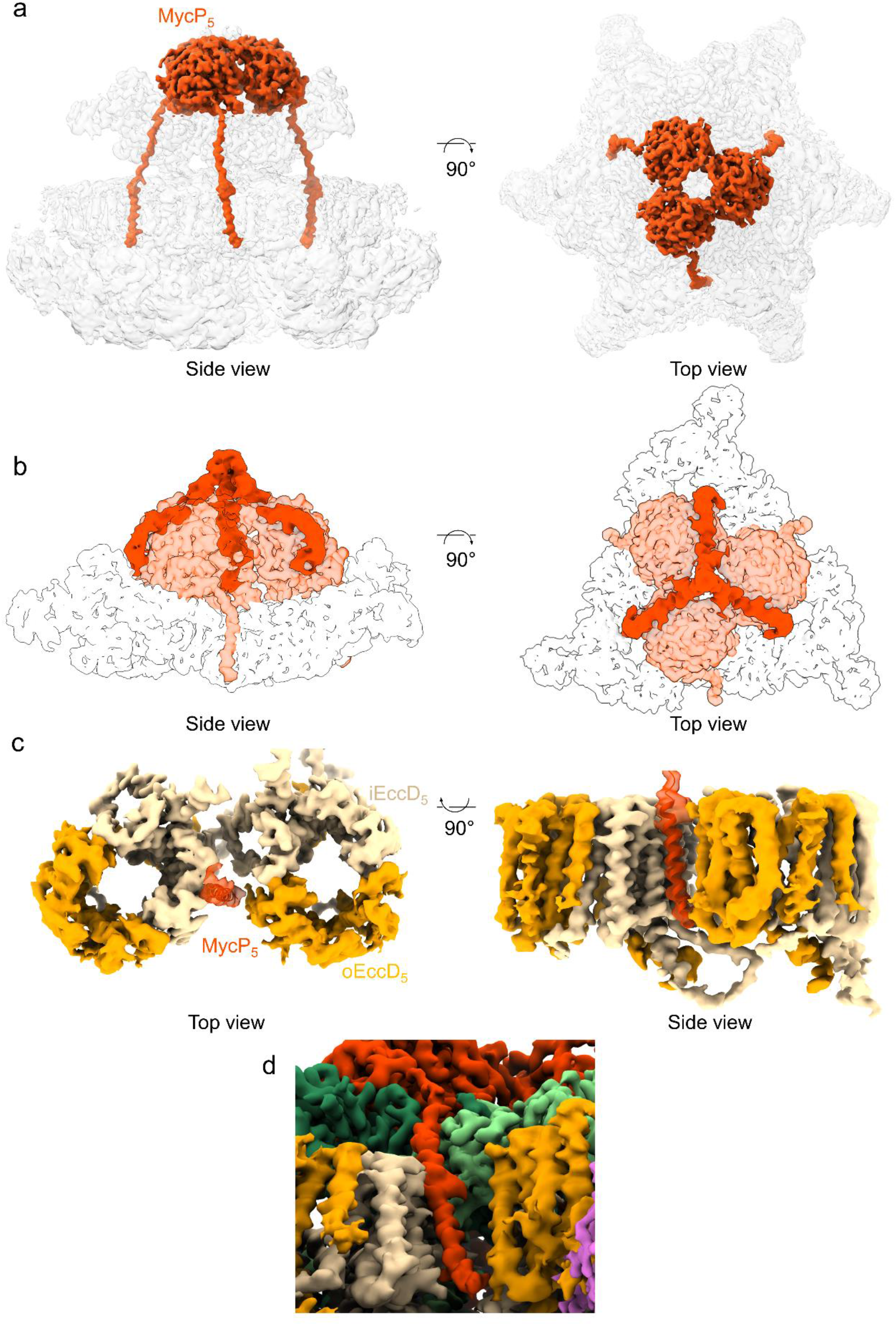
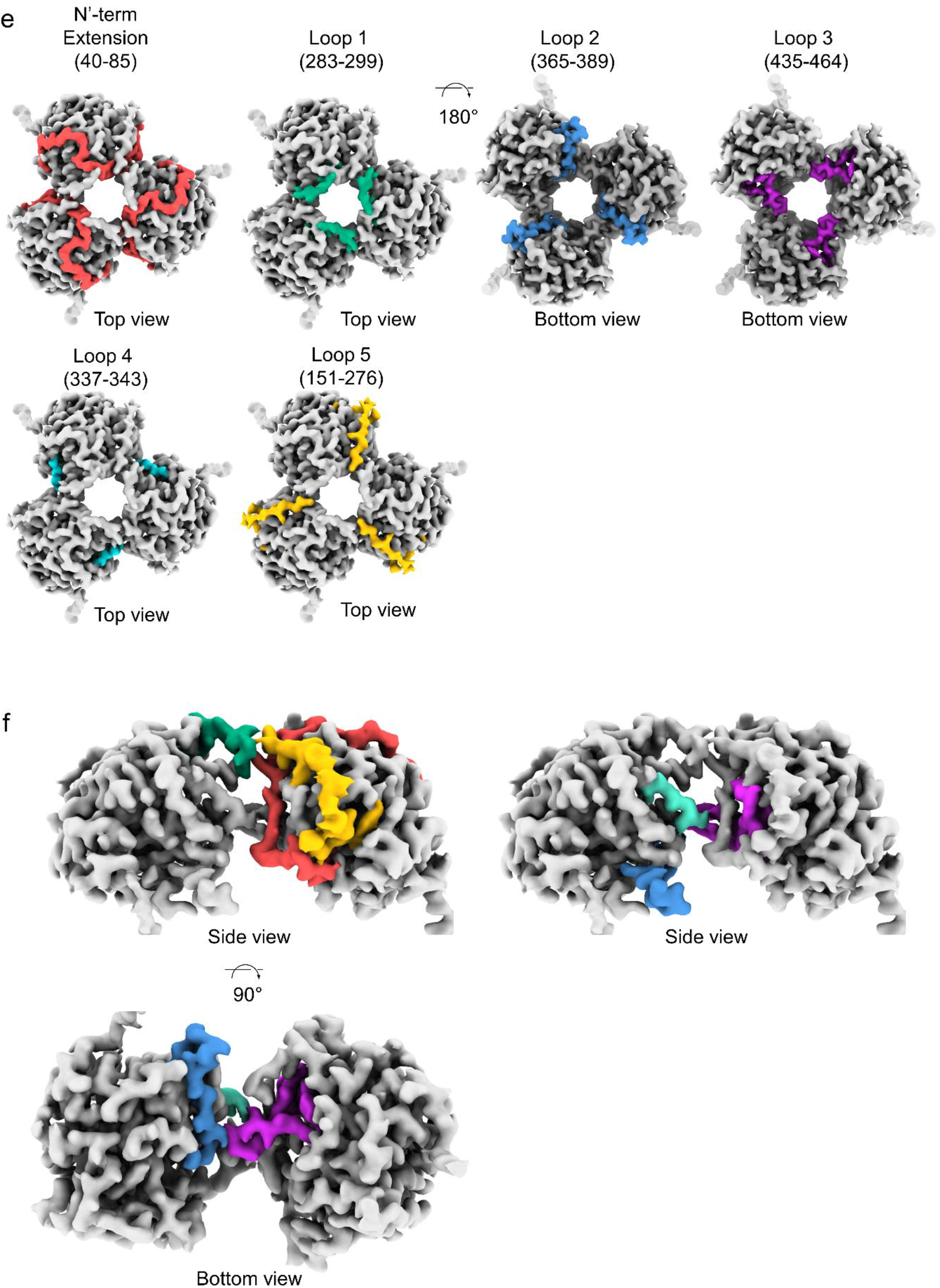
MycP_5_ caps a periplasmic cavity with its active site directed towards the lumen. **a**, Side and top view of an intact ESX-5_mtb_ assembly with MycP_5_ coloured as in Fig. 1 and the rest of the components transparent. **b**, Side and top view of the periplasmic assembly with EccB_5_ in white, the MycP_5_ density shown in transparent red and loop 5 of MycP_5_ depicted in solid red. Loop 5 folds along the protease domain, towards the pore formed by the MycP_5_ trimer. At higher thresholds, loop 5 caps this pore. **c**, Top and side view of a dimer of EccD_5_ barrels, of which one barrel (left) binds via iEccD_5_ to the MycP_5_ TMH. **d**, Side bottom view of an ESX-5 dimer, centred at the TMH of MycP_5_, highlighting the interaction of the TMH with iEccD_5_ at the periplasmic side of the inner membrane. **e**, Top or bottom view of MycP_5_ trimers depicted in grey with the domains that are involved in MycP_5_:MycP_5_ interactions depicted in different colours. **f**, Side views showing the MycP_5_:MycP_5_ interactions mediated by the same domains depicted in the same colours as in e.

**Extended Data Fig. 11.**
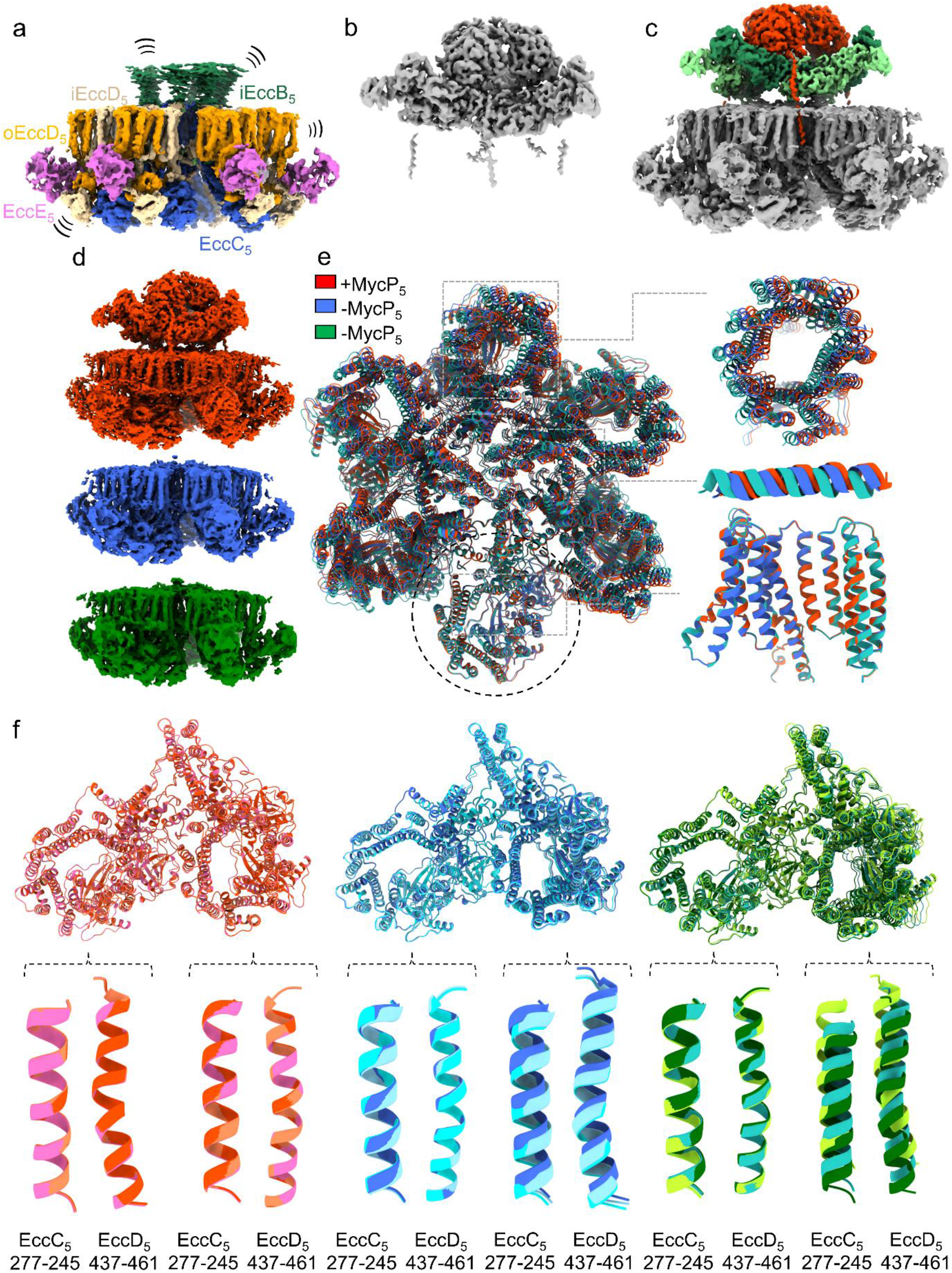
MycP_5_ acts as an allosteric driver for periplasmic EccB_5_ hexamerization and complex stability. **a**, Cryo-EM density map of a MycP_5_-free ESX-5_mtb_ membrane complex, zoned and coloured as in Fig. In the absence of MycP_5_, the periplasmic domains of EccB_5_ display high flexibility. The rest of the membrane complex displays increased heterogeneity when compared to the MycP_5_-bound map. **b**, Map of difference created by subtracting the MycP_5_-free map from the MycP_5_-bound map. **c**, Overlay of a and b. **d**, MycP_5_-bound map in red and the two MycP_5_-free maps in blue and green. **e**, A model of the MycP_5_-bound map, in which MycP_5_ and residue 84-504 of EccB_5_ were removed, was fitted into the models of the two MycP_5_-free maps, as specified in Material and Methods. Models were aligned at one EccD_5_ barrel (dark dotted circle), revealing significant variations and shifts between the three maps. Upper inset shows that there is consistent variation between all three maps at membrane level (EccD_5_ barrel); Middle inset shows variations between maps at cytosolic level (EccB_5_ N-terminal helix, 20-38); Lower inset highlights iEccD_5_ from the EccD_5_ barrel that was used for the alignment, showing that overall protomer structure does not change in the absence of MycP_5_. **f**, Dimers from every individual map, color-coded the same as in d, were extracted and aligned to each other on one EccD_5_ barrel (left) as in e. Insets from these alignments, derived from both protomers, show that all three dimers of the MycP_5_-bound map show little to no variation, whereas the two MycP_5_-free maps show a higher degree of heterogeneity between dimers.

**Extended Data Fig. 12.**
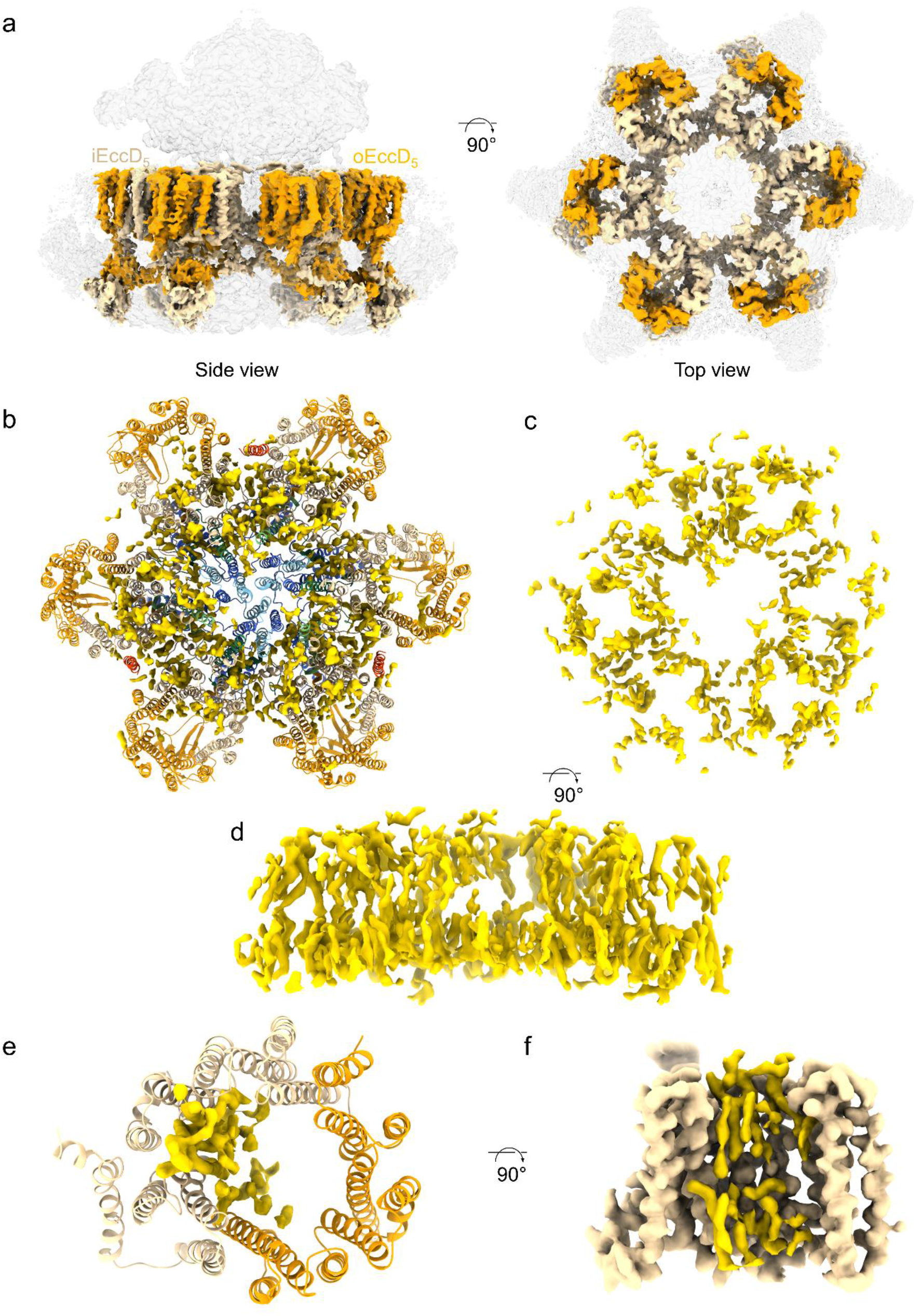
Six lipid-filled EccD_5_ barrels form a central raft. **a**, Side and top view of an intact ESX-5_mtb_ assembly, in which i- and oEccD_5_ is coloured as in Fig. 1 and the rest of the components are transparent. **b**, Top view of the membrane region of the ESX-5_mtb_ model overlayed with observed lipids, coloured in bright yellow. **c**, Same view as b, but only showing the lipids. **d**, Same image as c, but rotated 90 degrees to show a side view, highlighting a bilayer-like lipid structure. **e**, Top view of an EccD_5_ barrel with observed lipids bound to iEccD_5_. **f**, Side view of an iEccD_5_ monomer overlaid with observed lipids.

**Extended Data Fig. 13.**
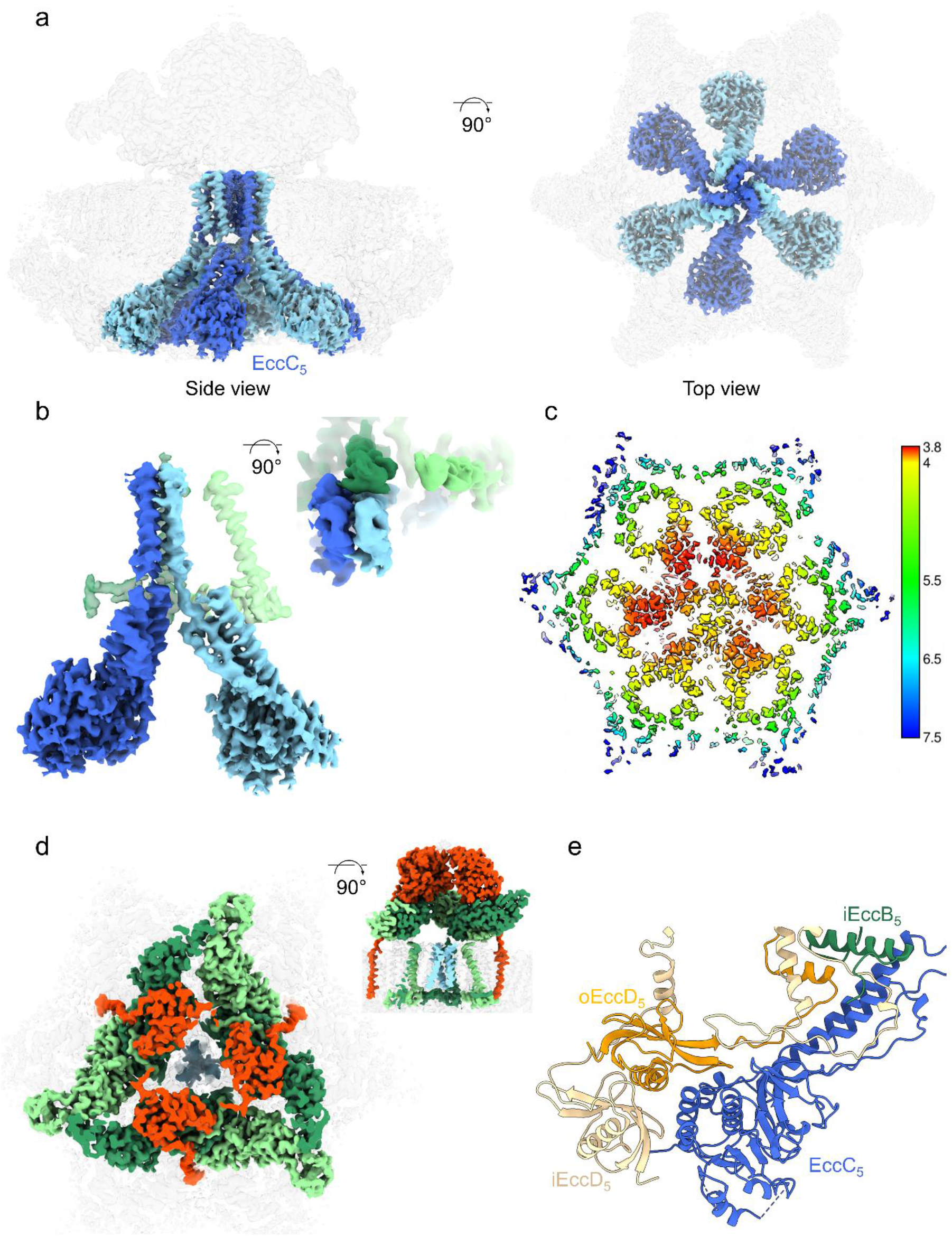
Three four TMH-bundles of EccC_5_ gate a central pore. **a**, Side and top view of an intact ESX-5_mtb_ assembly, in which EccC_5_ is coloured in alternating light and dark blue and the rest of the components are transparent. **b**, Extracted dimeric EccC_5_ and the TMHs and N-termini of EccB_5_ from the same dimer. 90 degrees inset rotation shows that the TMH of oEccB_5_ is not contacted by the TMHs of EccC_5_. **c**, Top membrane cross section through a local resolution map, displaying decreased resolution of the central space occupied by the TMHs of EccC_5_, compared to the surrounding EccB_5_ basket and TMHs of iEccD_5_. **d**, Top view of the full membrane complex with the EccC_5_ TMH-pyramid in light blue and the EccB_5_:MycP_5_ periplasmic cavity in the same colours as in Fig.1. Shown is that the EccC_5_ TMH-pyramid aligns with the periplasmic cavity and the MycP_5_ formed pore. MycP_5_ top part is partially sectioned, for clarity. Inset showing a 90 degrees rotation side cross section of the same map. **e**, Ribbon model highlighting the structural features of the cytosolic bridge.

**Extended Data Fig. 14.**
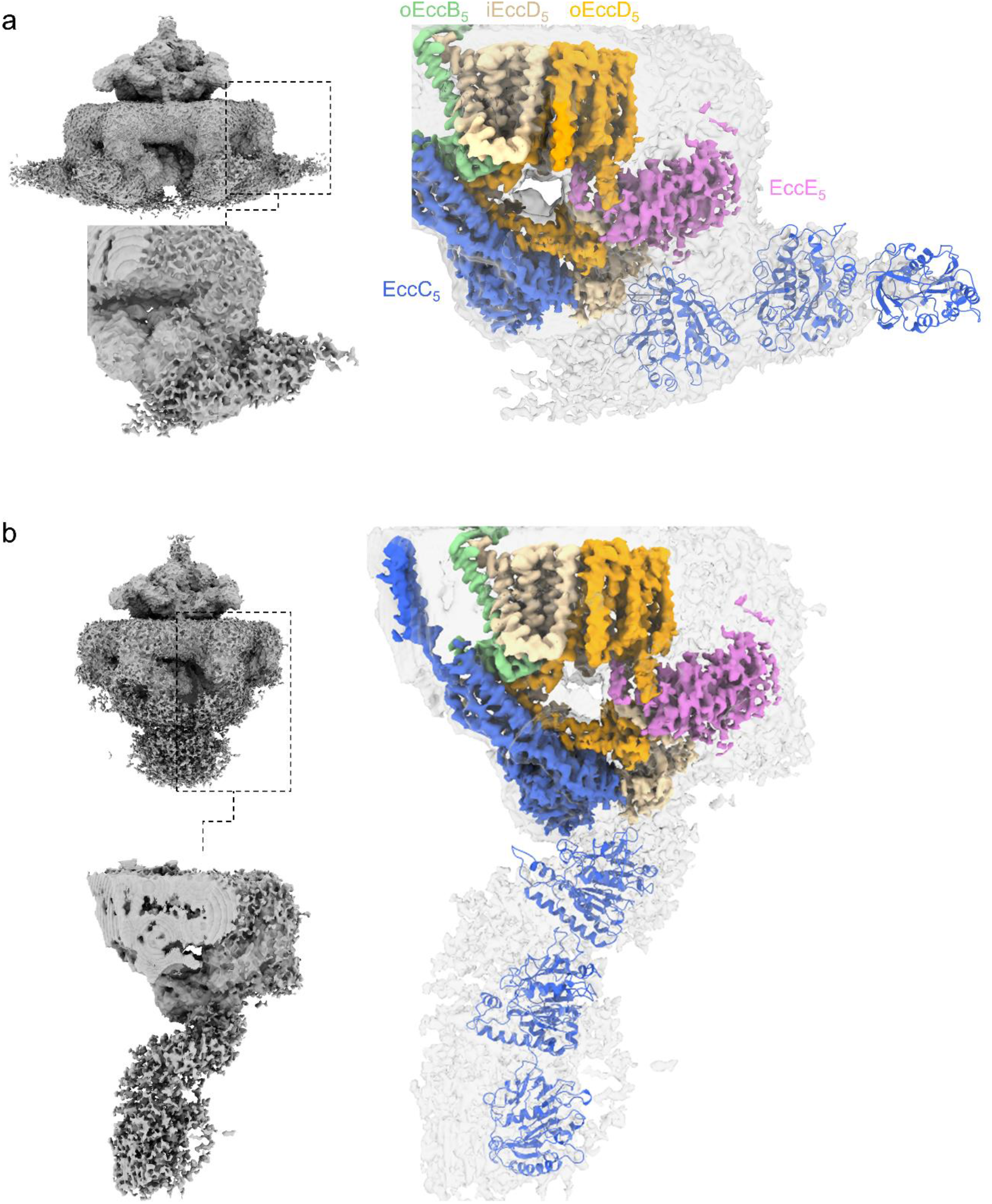
EccC_5_ adopts two separate conformations. **a**, Extended conformation in which an EccC_5_ NBD1-3 model is fitted to highlight the overall position of these domains with respect to the rest of the membrane complex. **b**, Similar as in a, but for the contracted conformation.

**Extended Data Fig. 15.**
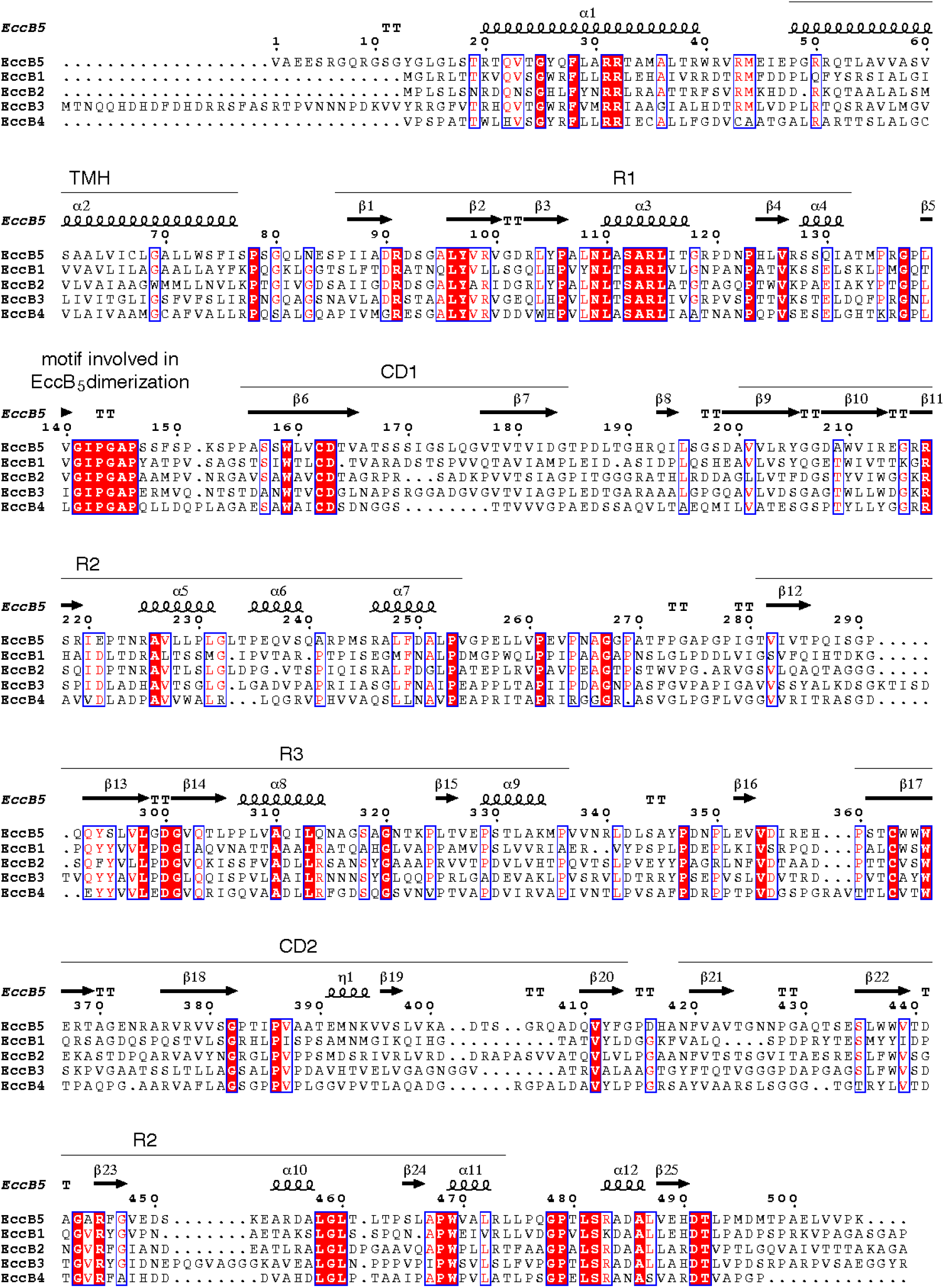
Structure based sequence alignment of the five EccB paralogs of *M. tuberculosis* H37Rv. Numbering and secondary structure elements are derived from the EccB_5_ sequence and structure. Sequences coding for the TMH, domain R1, R2, R3, R4 and two sequences (CD1 and CD2) that code for the central domain (CD) are indicated by black lines. The conserved GIPGAP motif involved in EccB_5_ dimerization is highlighted. The sequence alignment is produced using ESPript (http://espript.ibcp.fr)^17^.

**Extended Data Fig. 16.**
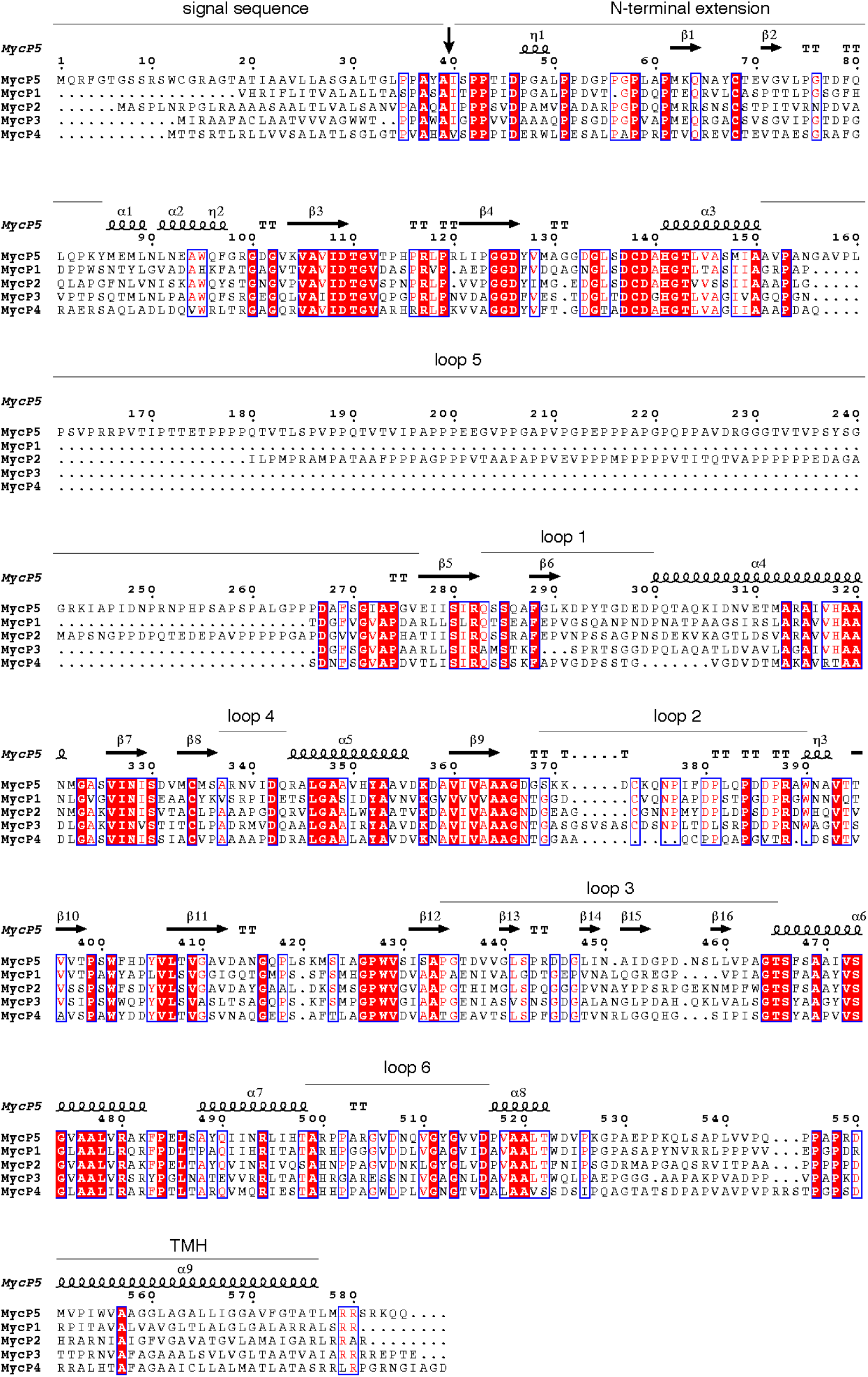
Structure based sequence alignment of the five MycP paralogs of *M. tuberculosis* H37Rv. Numbering and secondary structure elements are derived from the MycP_5_ sequence and structure. The signal protease cleavage site is indicated by a vertical arrow. Sequences coding for the signal sequence, N-terminal extension, loops 1 to 6 and the TMH are indicated by a black lines. While loop 1, 2, 3 and 5 have been assigned previously^18,19^, loop 4 and 6 are newly assigned in this study. The sequence alignment is produced using ESPript (http://espript.ibcp.fr)^17^.

**Extended Data Table 1.**
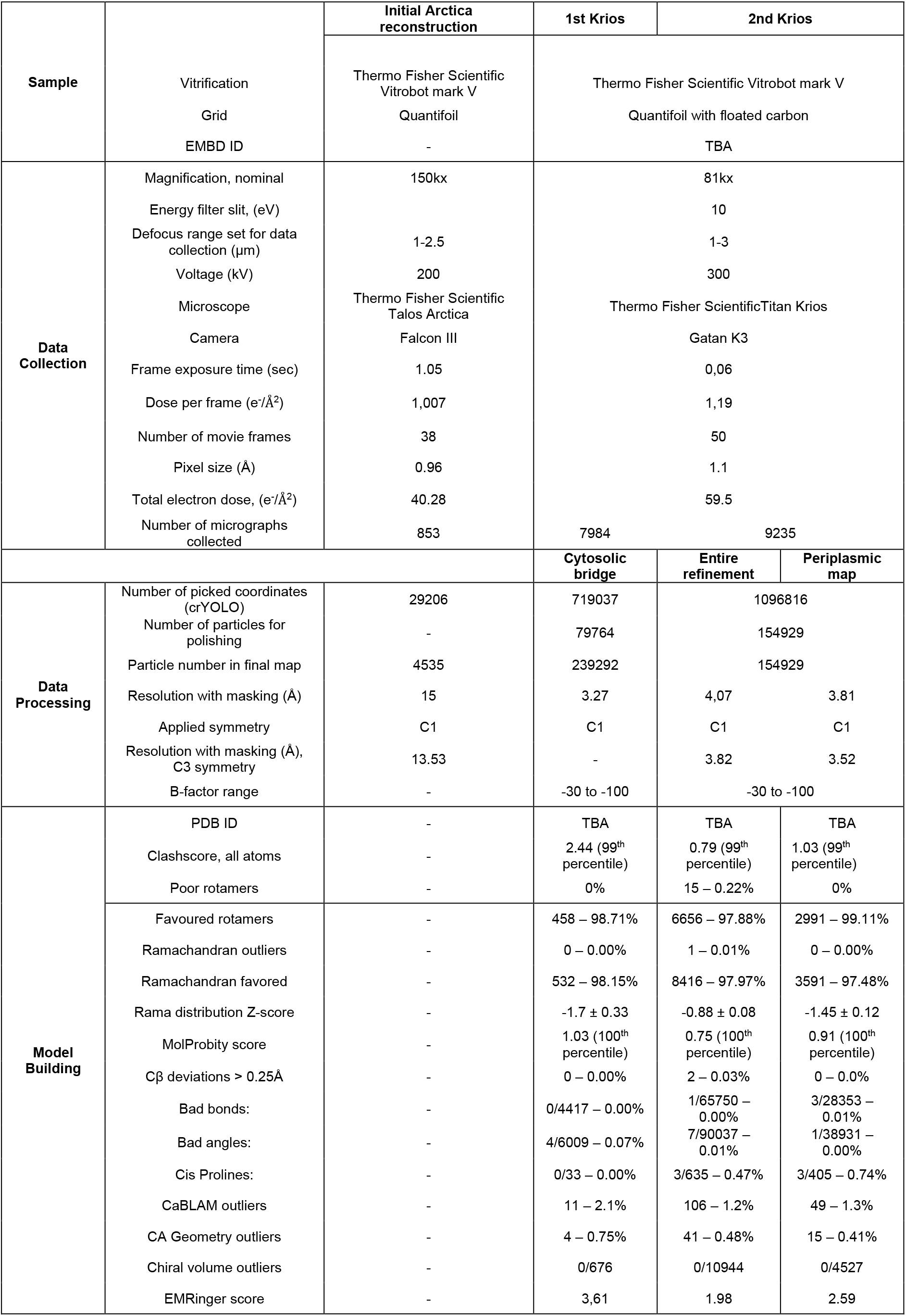
Cryo-EM data collection, processing and model building summary.

**Extended Data Table 2.**
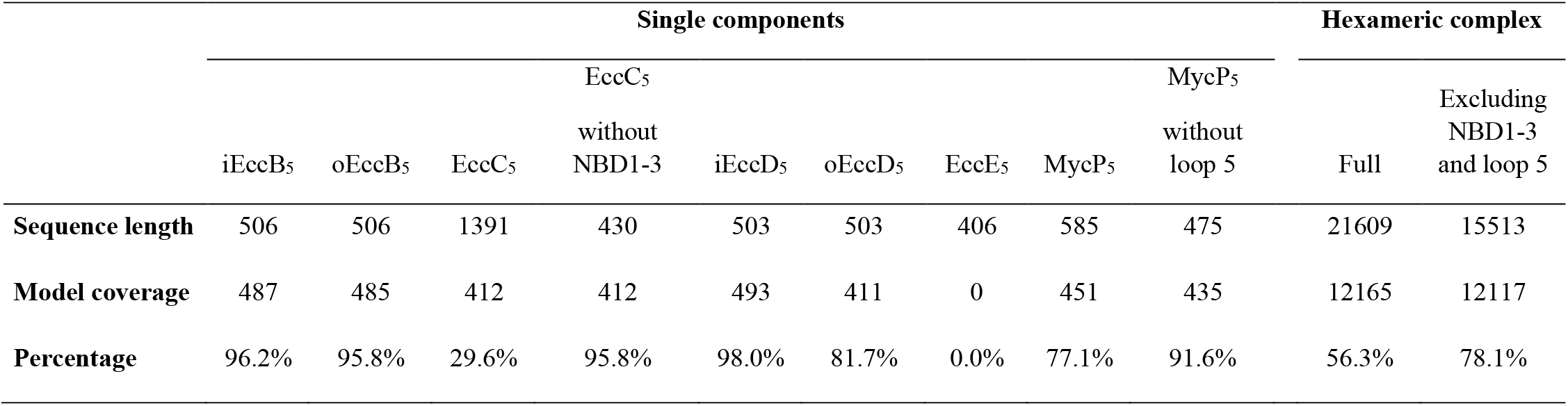
Model sequence coverage.

**Extended Data Table 3.**
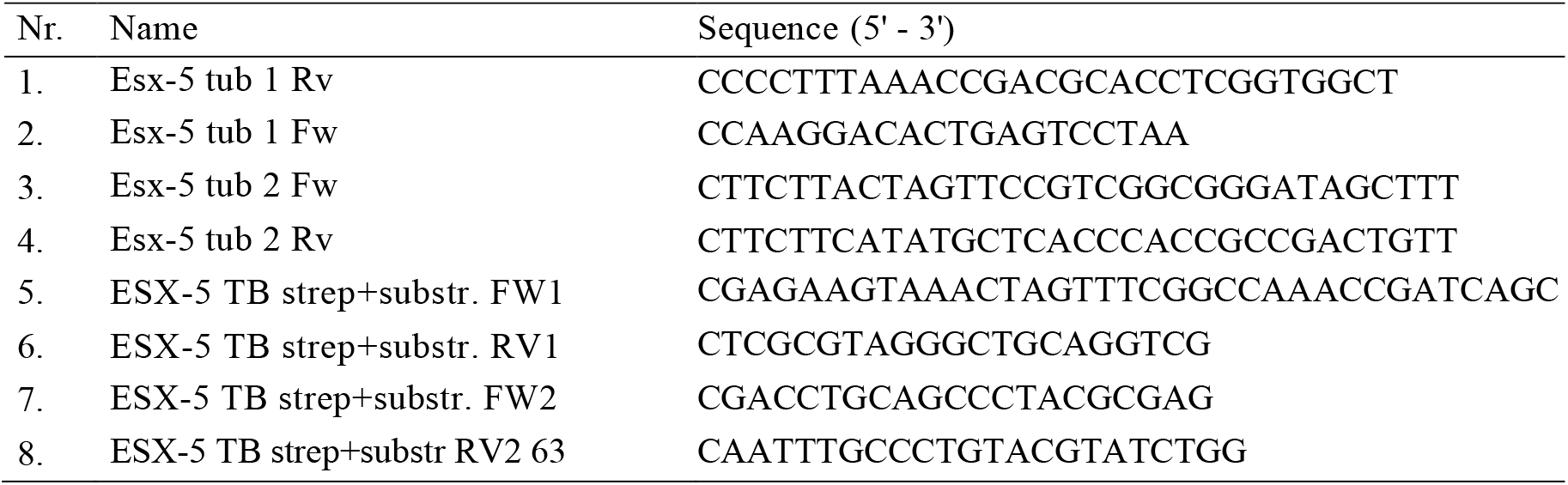
List of primers used in this study.

---

Extended Data Movie 1. Overall structure of the ESX-5_mtb_ membrane complex

Extended Data Movie 2. The periplasmic dome of the ESX-5_mtb_ membrane complex

Extended Data Movie 3. The central, EccC_5_ gated, pore of the ESX-5_mtb_ membrane complex

Extended Data Movie 4. Flexibility of the MycP_5_ free maps and stabilization upon MycP_5_ binding.

## Notes

### Competing Interest Statement

The authors have declared no competing interest.

